# Biomolecular condensation of ERC1 recruits ATG8 and NBR1 to drive autophagosome formation for plant heat tolerance

**DOI:** 10.1101/2024.09.09.611939

**Authors:** Ka Kit Chung, Ziwei Zhao, Kai Ching Law, Juncai Ma, Cheuk Him Chiang, Kwan Ho Leung, Ruben Shrestha, Yixin Wu, Chaorui Li, Ka Ming Lee, Lei Feng, Xibao Li, Kam Bo Wong, Shou-Ling Xu, Caiji Gao, Xiaohong Zhuang

## Abstract

Macroautophagy (hereafter autophagy) is essential for cells to respond to nutrient stress by delivering cytosolic contents to vacuoles for degradation via the formation of a multi-layer vesicle named autophagosome. A set of autophagy-related (ATG) regulators are recruited to the phagophore assembly site for the initiation of phagophore, as well as its expansion and closure and subsequent delivery into the vacuole. However, it remains elusive that how the phagophore assembly is regulated under different stress conditions. Here, we described an unknown *Arabidopsis (Arabidopsis thaliana)* cytosolic ATG8-interaction protein family (ERC1/2), that binds ATG8 and NBR1 to promote autophagy. ERC1 proteins translocate to the phagophore membrane and develop into classical ring-like autophagosomes upon autophagic induction. However, ERC1 proteins form large droplets together with ATG8e proteins when in the absence of ATG8 lipidation activity. We described the property of these structures as phase-separated membraneless condensates by solving the *in vivo* organization with spatial and temporal resolution. Moreover, ERC1 condensates elicits a strong recruitment of the autophagic receptor NBR1. Loss of ERC1 suppressed NBR1 turnover and attenuated plant tolerance to heat stress condition. This work provides novel insights into the mechanical principle of phagophore initiation via an unreported ERC1-mediated biomolecular condensation for heat tolerance in *Arabidopsis*.

## Introduction

Autophagy is conservedly operated in most eukaryotic cells to eliminate the detrimental or dysfunctional components, which is highly induced upon stress conditions (Liu and Bassham, 2012; Marshall and Vierstra, 2018; Isono et al., 2024; Otegui et al., 2024; Zhuang et al., 2024). A double-membrane compartment called autophagosome is formed to engulf and degrade the harmful materials during autophagy. During this process, multiple autophagy-related (ATG) proteins and other regulators are recruited for either autophagosome formation and/or cargo recognition. ATG8 family proteins, are conjugated on the autophagosome membrane and function throughout the whole autophagic process. In a general model for autophagosome development, ATG machinery is activated and recruited to the phagophore assembly site for the development of phagophore. Subsequently, ATG8 proteins are conjugated to the phagophore membrane through a ubiquitin-like conjugation process, which is essential for cargo recognition and autophagosome closure. Autophagosomes can be highly induced under starvation conditions to bulkily engulf cytoplasmic cargoes. Alternatively, specific cargoes (proteins or organelles) are recognized directly or indirectly by ATG8 proteins via selective autophagic receptors (SARs), thus leading to specific cargo sequestration into the phagophore, which is known as selective autophagy (Liu and Bassham, 2012; Isono et al., 2024; Otegui et al., 2024; Zhuang et al., 2024). Once the phagophore is sealed to become a complete autophagosome, it will be sent to the vacuole together with the contents or undergo further maturation process via fusion with the endosomes.

A number of novel autophagic regulators containing ATG8-family interacting motif (AIM) or LC3-interacting region (LIR) have been discovered and extensively studied in yeast and mammals (Kirkin and Rogov, 2019). These ATG8-binding proteins either coordinate with the ATG machinery to orchestrate autophagosome formation, or to exert selectivity as SARs in selective autophagy, underscoring the complexity in autophagy regulation network governing cellular homeostasis in response to different stress conditions. NBR1 represents one of the extensively studied SAR in plants that binds to ATG8 in a classical AIM-dependent manner, which has been implicated to function in multiple types of selective autophagy (Svenning et al., 2011; Zhou et al., 2013; Hafren et al., 2017; Jung et al., 2020a; Thirumalaikumar et al., 2021; Lee et al., 2023; Ebstrup et al., 2024). In recent years, advances in proteomic analysis have allowed researchers to identify several novel plant ATG8-interacting proteins for different types of selective autophagy, like C53 in ERphagy, TraB in mitophagy, and clathrin in Golgiphagy (Stephani et al., 2020; Li et al., 2022; Zhou et al., 2023). These exciting achievements have greatly advanced our understanding of selective autophagy in plants for different stress responses. Meanwhile, advances in studying ATG8-interacting regulators have also shed new lights into the underlying molecular mechanisms for autophagosome formation, ranging from phagophore initiation, autophagosome maturation and vacuolar delivery (Sutipatanasomboon et al., 2017; Kim et al., 2022; Sun et al., 2022; Zhao et al., 2022; Kim et al., 2023; Zeng et al., 2023; Mosesso et al., 2024). Nevertheless, ATG8-binding regulators for autophagosome initiation are largely unexplored, and the molecular property and regulation mechanism underlying phagophore initiation remain elusive in plants.

Using a proximity labeling approach with *Arabidopsis* ATG8e tagged with TurboID as a bait, we found a protein family with unknown function, which contains two homologs (AT4G02880 and AT1G03290). They both contain coil-coil domains, which are predicted to closely relate to the mammalian ELKS/Rab6-interacting/CAST (ERC) protein family, a central scaffold that forms a complex of multidomain proteins to mediate synaptic vesicle secretion (Monier et al., 2002; Ohtsuka et al., 2002; Wang et al., 2002). As there is no available information about this protein family in plants, we hence named these two proteins as ERC1 and ERC2 respectively. Interaction mapping demonstrated a physical interaction between ERC1 and ATG8e through a C-terminus AIM-like motif of ERC1. We further showed that ERC1 responds to autophagic induction and localizes on autophagosome membrane. Loss of ATG8 conjugation activity induces overaccumulation of ERC1 together with ATG8e on large spherical condensates. Using correlative-light electron microscopy (CLEM), immuno-electron microscopy (EM), 3D electron tomography (ET) and dynamic analysis, we elucidated the liquid-like property of ERC1 condensate and its morphological features, unveiling a complex spatial architecture of the ERC1 condensate and its close association with the ER membranes. We further demonstrated that ERC1 also interacts with the autophagic receptor NBR1. Loss of ERC1 suppresses NBR1 turnover and heat stress resistance in *Arabidopsis*. Altogether, these data implicate ERC1 as a novel regulator via biomolecular condensation with ATG8 and NBR1 to drive autophagosome formation for plant stress resistance.

## Results

### *Arabidopsis* ERC protein family associate with ATG8 protein family

Our previous study has reported that dysfunction of ATG2 blocks autophagosome formation, resulting accumulation of unclosed autophagosomal structures (Luo et al., 2023). To search for novel ATG8-associated proteins on the unclosed autophagosomal structures, we generated *Arabidopsis* plants expressing TurboID-ATG8e in *atg2-1* mutant and conducted a proximity labeling analysis. From the mass spectrometry result, we found a protein family with unknown function, which contains two homologs (AT4G02880 and AT1G03290) (**Supplementary Figure S1A**). Both ERC1 and ERC2 contain conserved coil-coil domains within the N- and C- terminuses, as well as two intrinsically disordered regions (IDRs) (**Figure 1A**). As a first step to characterize the ERC protein family in *Arabidopsis*, we performed a phylogenetic tree to understand the evolution of the ERC protein family in plants (**Figure 1B**). Intriguingly, ERC protein family represents an uncharacterized conserved coil-coil domain containing family which can be found in 12 representative species in land plants. In addition, most of them also share high similarity in the N-terminus and C-terminus regions. Next, we expressed the GFP-tagged ERC1 or ERC2 with mCherry-ATG8e to examine their subcellular association in *Arabidopsis* protoplast cells. Both ERC1-GFP and ERC2-GFP fusion proteins colocalized with mCherry-ATG8e (**Figure 1C**). We also performed a recruitment assay in *Arabidopsis* protoplasts using an ER-anchored CNX-RFP-ATG8e chimera (Sun et al., 2022) with ERC1-GFP or ERC2-GFP. As shown in **Figure 1D and 1E**, CNX-RFP-ATG8e is well overlapped with ERC1-GFP and ERC2-GFP respectively, but no obvious colocalization between the CNX-RFP and ERC1-GFP or ERC2-GFP is observed. To test whether the association is specific to ATG8e, we performed another recruitment assay using CNX-mCherry-ERC1 with the nine ATG8 isoforms (ATG8a-ATG8i). Our result showed that all ATG8 isoforms were colocalized with CNX-mCherry-ERC1. To further confirm the interaction, we performed co-immunoprecipitation (Co-IP) assays, showing that Flag-tagged ERC1 or ERC2 is associated with YFP-ATG8e when transiently expressed in *Arabidopsis* protoplasts (**Figure 1F and 1G**). We next carried out an *in-vitro* GST pull-down assay using purified GST-ATG8e and ERC1 proteins, showing that ERC1 displays a higher binding affinity with GST-ATG8e than GST (**Figure 1H**). In order to study the function of ERC protein family in *Arabidopsis* plants, we further checked the transcription profile of the ERC protein family in different *Arabidopsis* tissues. Our data showed that ERC1 is ubiquitously expressed in root, stem, leaf and flower, whereas ERC2 is barely detected in root, leaf and flower (**Figure 1I**). Therefore, we selected ERC1 for the following biochemical and cellular analysis in *Arabidopsis* plants.

**Figure 1.**
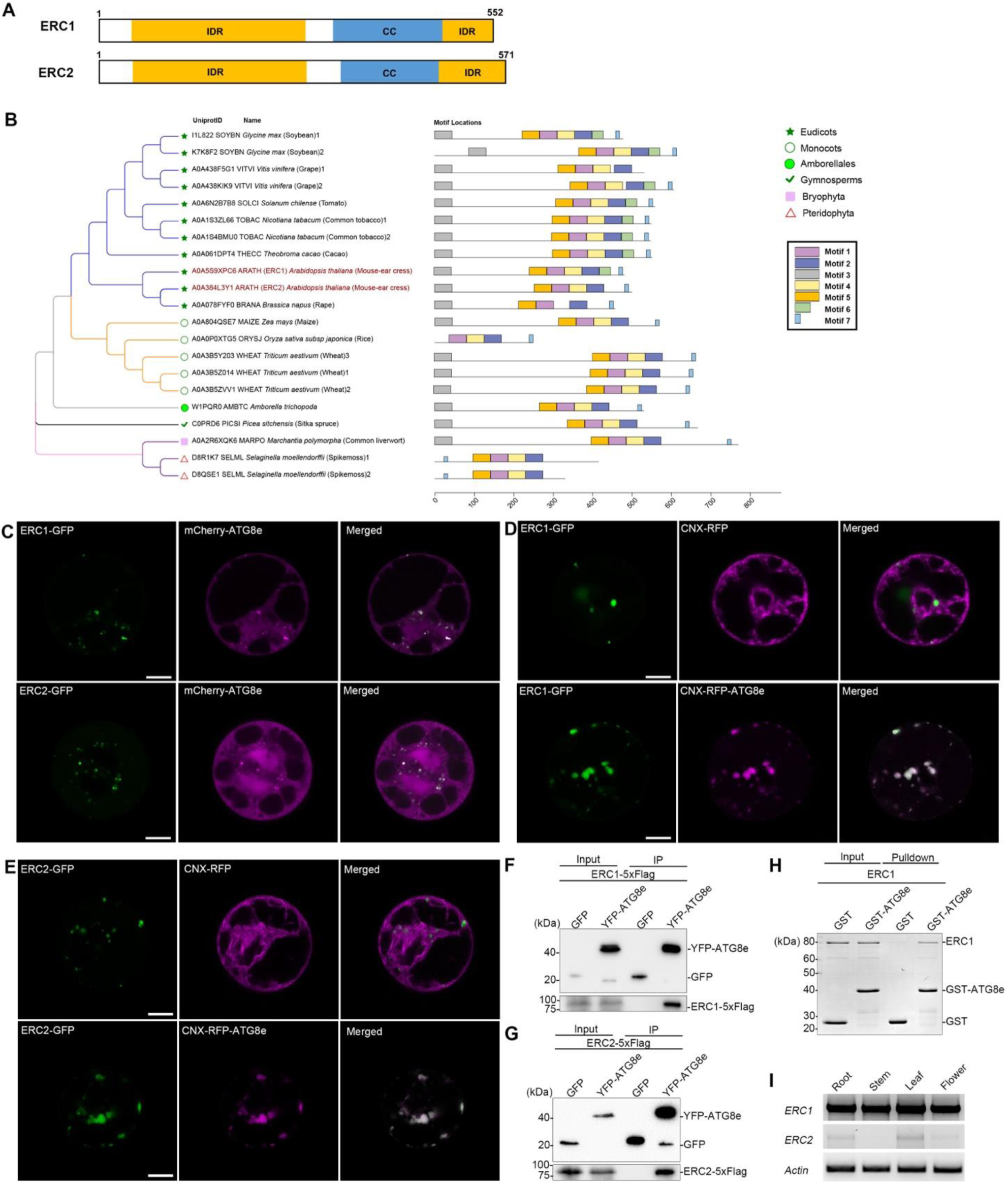
Arabidopsis ERC protein family associates with ATG8e. **(A)** Domain architectures of Arabidopsis ERC1 and ERC2. CC, coil-coil domain; IDR, disordered protein region. **(B)** Phylogenetic analysis of ERC orthologs (left) and motif conservation (right) among the representative plant species, with seven distinct motifs identified and illustrated in different colors. **(C)** Both GFP-tagged ERC1 and ERC2 colocalized with mCherry-ATG8e when transiently expressed in *Arabidopsis* PSBD protoplasts. Scale bar=10um. Similar results were obtained from three different independent experiments. (**D-E**) Both GFP-tagged ERC1 and ERC2 were recruited by CNX-RFP-ATG8e but not CNX-RFP when transient expressed in *Arabidopsis* PSBD protoplasts. Scale bar=10 μm. Similar results were obtained from three different independent experiments. (**F-G**) Immunoprecipitation (IP) analysis showed that both ERC1 and ERC2 associated with ATG8e. Cell lysate from Arabidopsis PSBD protoplasts transiently expressing ERC1-5xFlag or ERC2-5xFlag together with YFP-ATG8e or GFP were subjected to GFP-trap assay. Input and IP samples were detected with anti-GFP and anti-Flag antibodies respectively. Similar results were obtained from three different independent experiments. **(H)** ERC1 interacts with ATG8e. GST pull-down assays were performed with purified GST or GST-ATG8e with ERC1 proteins using Glutathione magnetic beads. 10% of input and pulldown fraction was loaded. The input and pulldown fractions were visualized by Coomassie Blue staining. Similar results were obtained from three different independent experiments. **(I)** RT-PCR analysis of ERC1 and ERC2 transcript levels in different Arabidopsis tissues. Actin was used an internal control. Similar results were obtained from three different independent experiments.

### ERC1 interacts with ATG8e via a C-terminus AIM-like motif

By far, most of the reported ATG8-interacting proteins are featured with a short peptide sequence named ATG8-family interacting motif (AIM), also known as LC3-interacting region (LIR), with a canonical sequence “W/F/Y-X-X-L/I/V” for binding to the LIR/AIM docking site (LDS) in ATG8 (Stephani and Dagdas, 2020; Rogov et al., 2023; Isono et al., 2024). Since ERC1 displays a physical interaction with ATG8e proteins, we first searched AIM-like motifs in ERC1. Based on the prediction (https://ilir.warwick.ac.uk)(Jacomin et al., 2016), we tested the two AIM-like motifs with high scores by introducing mutations in these AIM-like motifs and subjected to the recruitment assay. However, mutations in these two predicted AIM-like motifs did not disturb the overlapping between ERC1 fusion proteins and CNX-RFP-ATG8e (**Supplementary Figure S2A**). To further map the molecular determinant required for ERC1-ATG8 interaction, we generated several GFP-tagged ERC1 truncations (T1, T2 and T3) as depicted in **Figure 2A**, for recruitment assay with CNX-RFP-ATG8e. Surprisingly, we observed that CNX-RFP-ATG8e is almost fully overlapped with a C-terminus T2 truncation, but not with the N-terminus T1 truncation, which retained a cytosolic pattern (**Figure 2B**). However, when a short C-terminus T3 truncation was used, it greatly impaired the association with CNX-RFP-ATG8e. We further conducted a pull-down assay to validate their interactions with ATG8e (**Figure 2C)**. In agreement with the recruitment assay, His-tagged ATG8e interacts with ERC1 T2 only, but not with T1 or T3 truncation.

**Figure 2.**
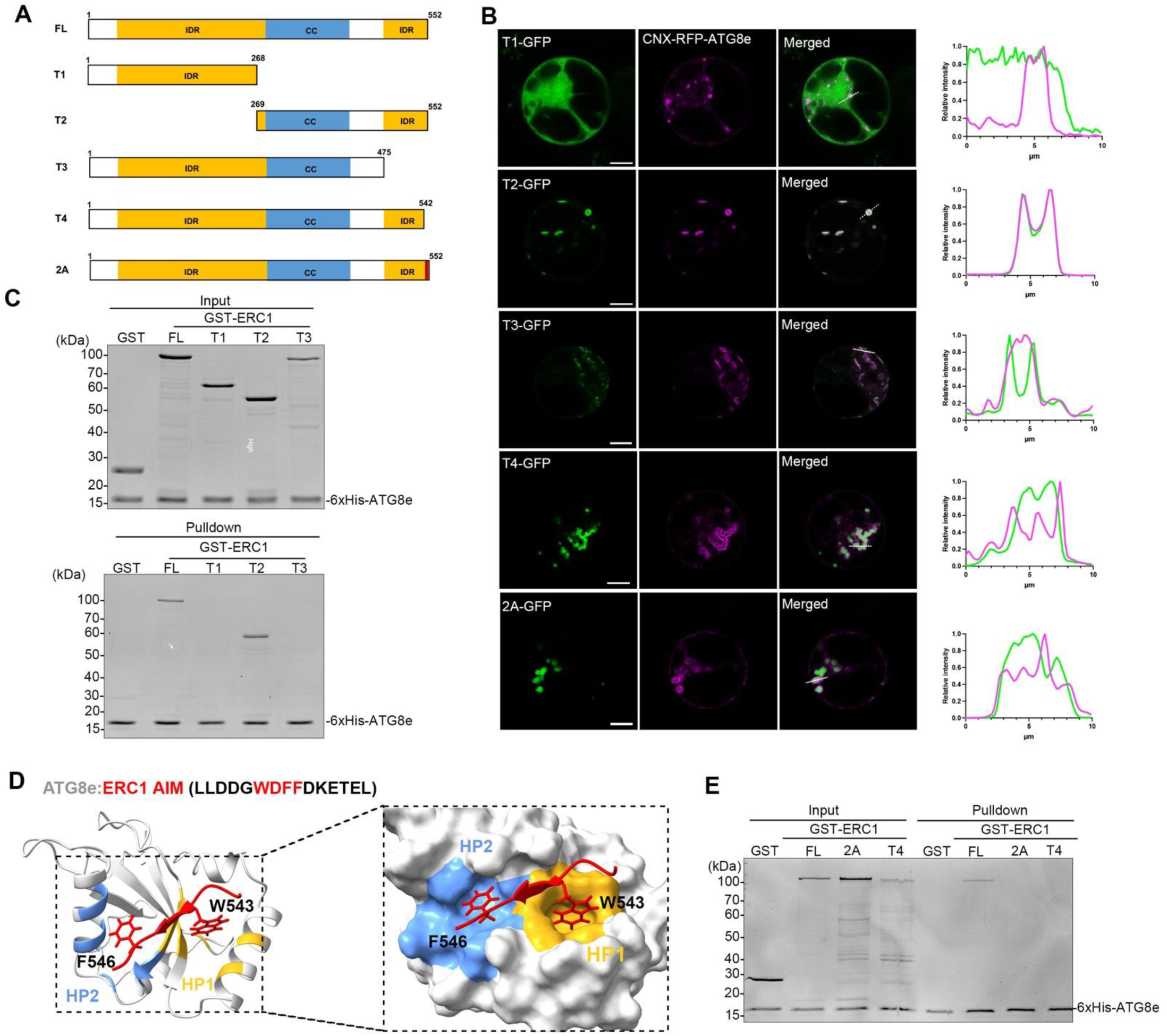
ERC1 interacts with ATG8e in an AIM-dependent manner. **(A)** Schematic of ERC1 full-length (FL) and deletion or mutation constructs (T1-T4, 2A); **(B)** Recruitment assay of ERC1 variants in (A) with CNX-RFP-ATG8e in transient expressed Arabidopsis protoplasts. CNX-RFP-ATG8e and GFP-tagged ERC1 variants were transiently co-expressed in Arabidopsis protoplasts and subjected to confocal observation. Scale bar= 10 μm. Fluorescence intensity profiles of GFP-tagged ERC1 variants and CNX-RFP-ATG8e for the indicated area labeled by the white dash lines were shown on the right. **(C)** His pull-down of 6xHis-ATG8e with GST or GST-tagged ERC1 truncations: T1, T2, T3. 10% of input and pulldown fraction was loaded. The input and pulldown fraction were visualized by Coomassie Blue staining. Similar results were obtained from three different independent experiments. **(D)** AlphaFold2-multimer predicts the AIM sequence of ERC1 docks into the ATG8e pockets, with the ERC1 AIM core residues “WDFF” in red and ATG8e backbone in grey. The side chains of ERC1 W543 and F546 identified by AF2 multimer for interacting with two hydrophobic pockets of ATG8e (HP1 in yellow and HP2 in blue) were highlighted. Area indicated by dashed box was enlarged and shown on the right. **(E)** His pull-down of 6xHis-ATG8e with GST or GST-tagged ERC1 truncations/mutants: FL, 2A, T4, 10% of input and pulldown fraction was loaded. The input and pulldown fraction were visualized by Coomassie Blue staining. Similar results were obtained from three different independent experiments.

These results prompted us to generate more truncations by chopping the C-terminus. Interestingly, removal of last 10 amino acids (“WDFFDKETEL”) in the C-terminus (T4) disturbed the association between ERC1 and ATG8e (**Figure 2B).** However, the “WDFFDKETEL” sequence did not match the feature of the canonical AIM sequences and was not shown in the list of online predicted AIM-like motifs. To evaluate whether it will form complex with ATG8e, we firstly used AlphaFold-multimer for protein complex prediction based on the last 15 amino acids of ERC1 and ATG8e (**Figure 2D**). Of note, AlphaFold-predicted model outlines a short motif of ERC1 (“WDFF”) is well accommodated in the LDS of ATG8e, with “W” and “F” showing close contacts with the two hydrophobic pockets (HP1 and HP2) respectively (**Figure 2D**). To further test the possible role of this sequence in ERC1-ATG8e interaction, we generated a mutation by alanine substitution of “WDFF” into “ADFA” (depicted as 2A) for pull-down and recruitment analysis (**Figure 2B and 2E**). In agreement with the predicted model, ERC1 2A mutation indeed compromised its association with ATG8e. Taken together, ERC1 interacts with ATG8 through a cryptic AIM motif.

### Subcellular localization of ERC1 in *Arabidopsis* plants

Next, we generated transgenic plants expressing ERC1-GFP, and crossed with different organelle markers to further characterize the subcellular behavior of ERC1 under different conditions. As shown in **Figure 3A**, under normal condition, ERC1-GFP mainly displayed a punctate pattern, which was partially overlapped with mCherry-ATG8e. We then subjected the seedlings to autophagic induction using Acibenzolar-S-methyl (BTH) treatment (Zhuang et al., 2013; Zhuang et al., 2017). As shown in **Figure 3A**, number of punctate labeled by ERC1-GFP were significantly upregulated, and some displayed cup-shaped or ring-like structures with different degrees of overlapping with mCherry-ATG8e, suggesting ERC1 might participate in different steps during autophagosome formation.

**Figure 3.**
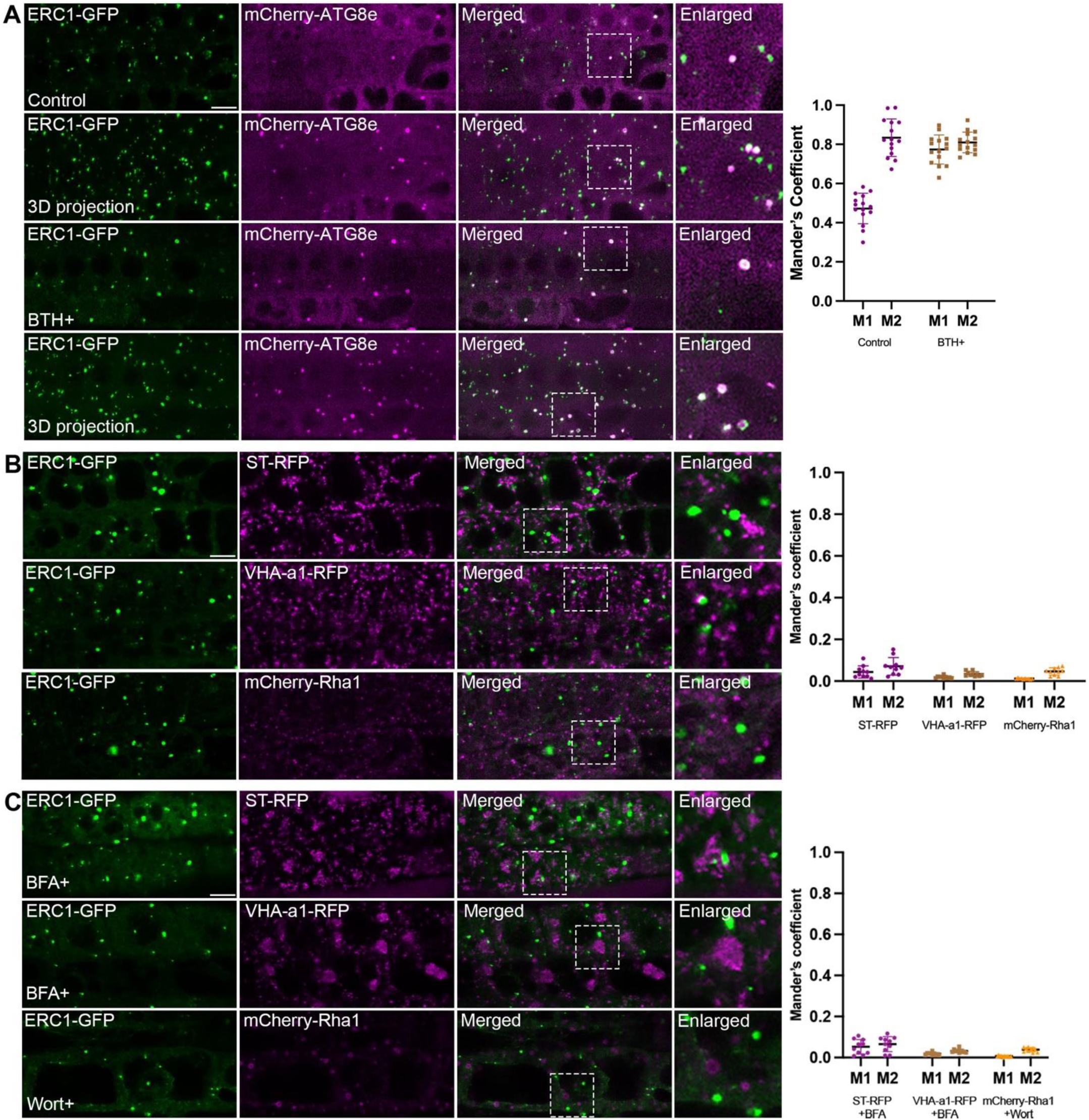
Subcellular localization of ERC1 in Arabidopsis root cells. **(A)** 5-day-old transgenic plants expressing ERC1-GFP/mCherry-ATG8e proteins were subjected to untreated or treated with BTH for 6 hours before imaging by confocal microscopy. Scale bar=10 μm. Quantification of the Mander’s colocalization coefficients between ERC1-GFP and the mCherry-ATG8e was shown on the right. M1, fraction of ERC1-GFP signal that overlaps with mCherry-ATG8e signal. M2, fraction of mCherry-ATG8e signal that overlaps with ERC1-GFP signal. Bars indicate the mean ± SD of 10 replicates. **(B)** 5-day-old transgenic plants expressing ERC1-GFP with Golgi marker (ST-RFP), TGN marker (VHA1-a1-RFP) or PVC/MVB marker (mCherry-Rha1) were subjected to untreated or treated with BFA or wortmannin before imaging by confocal microscopy. Quantification of the Mander’s colocalization coefficients between ERC1-GFP and the respective markers. M1, fraction of ERC1-GFP signal that overlaps with respective marker’s signal. M2, fraction of the respective marker’s signal that overlaps with ERC1-GFP signal. Bars indicate the mean ± SD of 10 replicates. **(C)** 5-day-old transgenic plants expressing ERC1-GFP with Golgi marker (ST-RFP), TGN marker (VHA1-a1-RFP) or PVC/MVB marker (mCherry-Rha1) were subjected to untreated or treated with BFA or wortmannin before imaging by confocal microscopy. Scale bar=10 μm. Quantification of the Mander’s colocalization coefficients between ERC1-GFP and the respective markers. M1, fraction of ERC1-GFP signal that overlaps with respective marker’s signal. M2, fraction of the respective marker’s signal that overlaps with ERC1-GFP signal. Bars indicate the mean ± SD of 10 replicates.

As majority of the ERC1-positive puncta are devoid of ATG8e signals under the normal condition, we hypothesized that ERC1 might resident on other subcellular compartments. Therefore, we crossed ERC1-GFP with Golgi marker (ST-RFP), TGN marker (VHA1-a1-RFP) or PVC/MVB marker (mCherry-Rha1) for confocal observation. However, we barely detected any overlapping between ERC1 and ST-RFP, VHA1-a1-RFP or mCherry-Rha1 (**Figure 3B)**. In addition, when we applied Brefeldin A (BFA), which causes the formation of BFA bodies by targeting the GTPase for Golgi and TGN trafficking (Geldner et al., 2003), it appears that the punctate pattern of ERC1-GFP remained unchanged (**Figure 3C**). Treatment of wortmannin (wort) that blocks PI3K activity and leads to enlarged ring-like MVB/PVC structures (Tse et al., 2004), ERC1-positive puncta remained unchanged and separated from ring-like structures labeled by mCherry-Rha1 (**Figure 3C**). These results indicate that the ERC1-GFP resides on distinct compartments that are separated from the endomembrane structures.

To gain more insights into the temporal relationship of ERC1 with ATG8 proteins during autophagosome formation, we performed real-time imaging analysis to monitor the dynamic behavior of ERC1-GFP x mCherry-ATG8e using a spinning disk confocal microscope. A representative montage was shown in **Figure 4A** and **Supplementary movie 1,** illustrating the development of an ERC1-GFP punctum into a complete ring-like structure. The ERC1-GFP signal is more condensate at the beginning as a bright dot when compared with that of mCherry-ATG8e. At around 19 min (frame 7), the signal of mCherry-ATG8e increased while the intensity of ERC1-GFP gradually weakened (**Figure 4B**), displaying a ring-like pattern overlapping with mCherry-ATG8e. This process lasts for around 10 min. At ∼ 40 min (frame 13), the ring-like structure is remodeled into a multi-ring structure, with an extending ring-like structure labeled by mCherry-ATG8e but devoid of ERC1-GFP (arrowhead).

**Figure 4.**
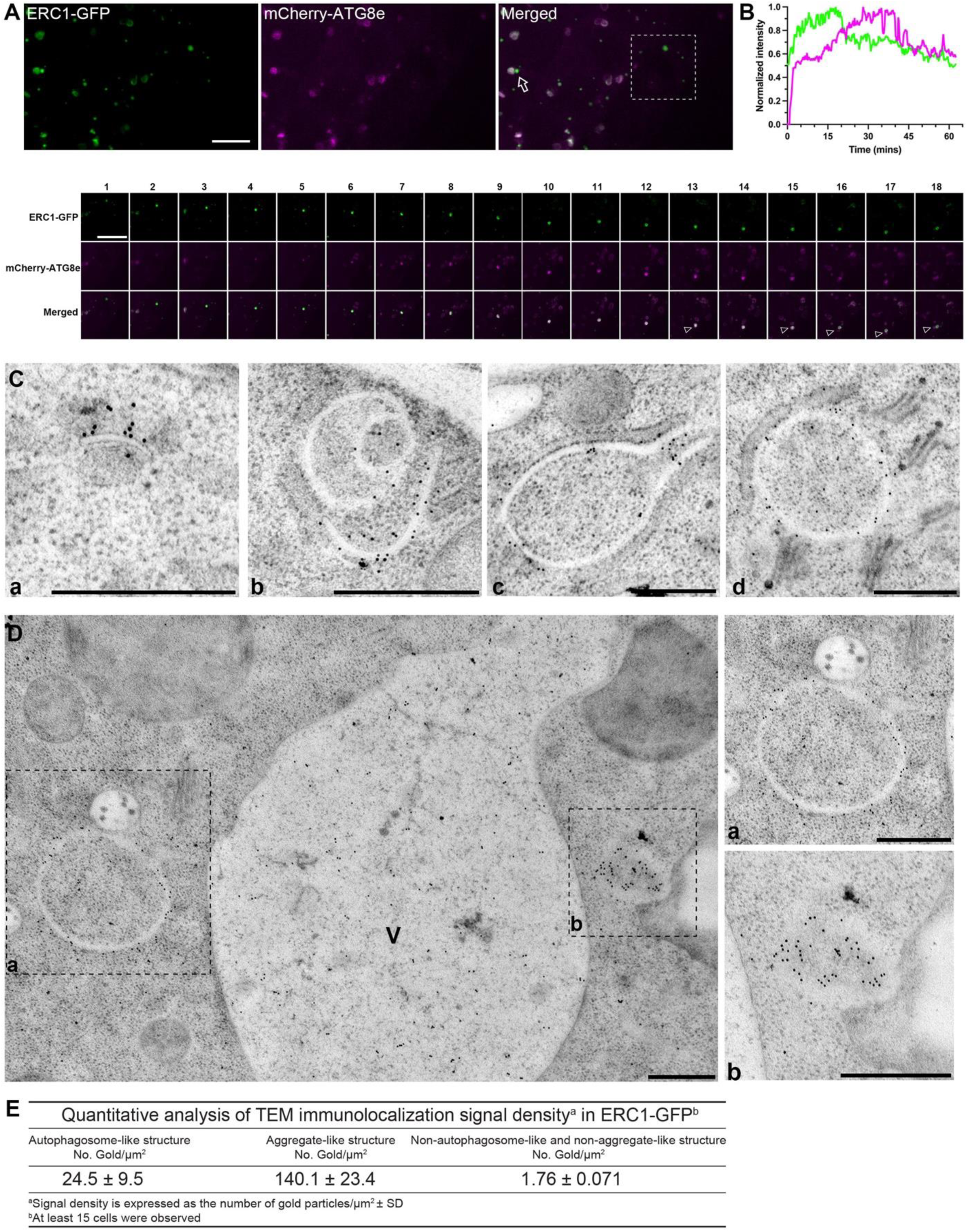
Real-time imaging and immune-gold electron microscopy analysis reveal ERC1-GFP proteins translocate to autophagosome membranes. **(A)** Real-time imaging showed the dynamic relationship between ERC1-GFP and mCherry-ATG8e. 5-days-old Arabidopsis seedlings expressing ERC1-GFP and mCherry-ATG8e with BTH treatment in ½ MS medium were subjected to observation under the spinning disk microscope. The dashed square indicated the cropped region for the time-lapse imaging analysis. Representative montages of the ERC1-GFP punctum and the mCherry-ATG8e structure are shown below. Scale bar, 5 µm. **(B)** Quantification of the normalized fluorescent intensities of ERC1-GFP (green) punctum and mCherry-ATG8e (magenta) punctum in the montage analysis from (A) by Fiji. Scale bar=5 μm. **(C)** Immuno-EM analysis reveals ERC1 proteins localized on autophagosomal membranes at different stages during autophagosome formation. 5-days-old root tips of ERC1-GFP with 6 hours BTH treatment in ½ MS medium were subjected to high-pressure freezing fixation. Sections were immunolabeled with anti-GFP antibodies, followed by gold particle labeling for anti-GFP (10 nm). Scale bar, 500 nm. **(D)** Immuno-EM analysis showed ERC1 proteins localize on a typical double membrane autophagosome (a) and a membraneless structure (b). **(E)** Quantitative analysis of gold-labeling density using the anti-GFP antibody against ERC1-GFP.

To further characterize these ERC1-GFP positive structures at the ultrastructure level, we conducted immuno-electron microscopy (EM) analysis. Root tips from 5-days-old transgenic plants expressing ERC1-GFP with BTH treatment were subjected to high pressure freezing (HPF) with freeze substitution, followed by gold labeling using GFP antibody against ERC1-GFP. Representative examples corresponding to different stages of autophagosome structures with gold labeling, including crescent-shape isolation membrane, unsealed multi-layered or double-layered structure, as well as typical complete double-layered autophagosome, were shown in **Figure 4C**. In addition, gold particles were enriched on the structure without a clear membrane boundary (**Figure 4D,b**), but displaying a unique density different from the surrounding or the lumen of autophagosome-like structure (**Figure 4D,a**).

### ERC1 forms large membraneless condensates together with ATG8e independent of the ATG8 conjugation system

Regarding that ERC1 was detected on phagophore membrane at a very early stage, we next exploited the possible relationship between ERC1 and other ATG machinery to understand how ERC1 participates in autophagy. Therefore, we introduced ERC1-GFP into several *atg* mutants, including *atg2-1* that impairs autophagosome closure, *atg5-1* and *atg7-2* which block ATG8 lipidation for autophagosome initiation (Thompson et al., 2005; Chung et al., 2010; Luo et al., 2023). Surprisingly, we observed signals of ERC1-GFP were hyperaccumulated on large droplet-like structures in the above *atg* mutants (**Supplementary Figure S3A**). Previous studies showed that fluorescent tagged ATG8 proteins mainly are localized in the cytosol with few large and bright puncta when lacking ATG5 or ATG7 activity, and overaccumulated on unclosed autophagosomes in the *atg2-1* mutant (Chung et al., 2010; Jung et al., 2020a; Luo et al., 2023). Strikingly, when we introduced both ERC1-GFP and mCherry-ATG8e into the *atg5-1* mutant, mCherry-ATG8e were also trapped on the large ERC1-GFP-positive structure (**Figure 5A and 5B**). Moreover, these droplet-like structures retained upon wortmannin or additional Conc A treatment (**Supplementary Figure S3B-C**), suggesting PI3K kinase activity is dispensable for the accumulation of these droplet-like structures.

**Figure 5.**
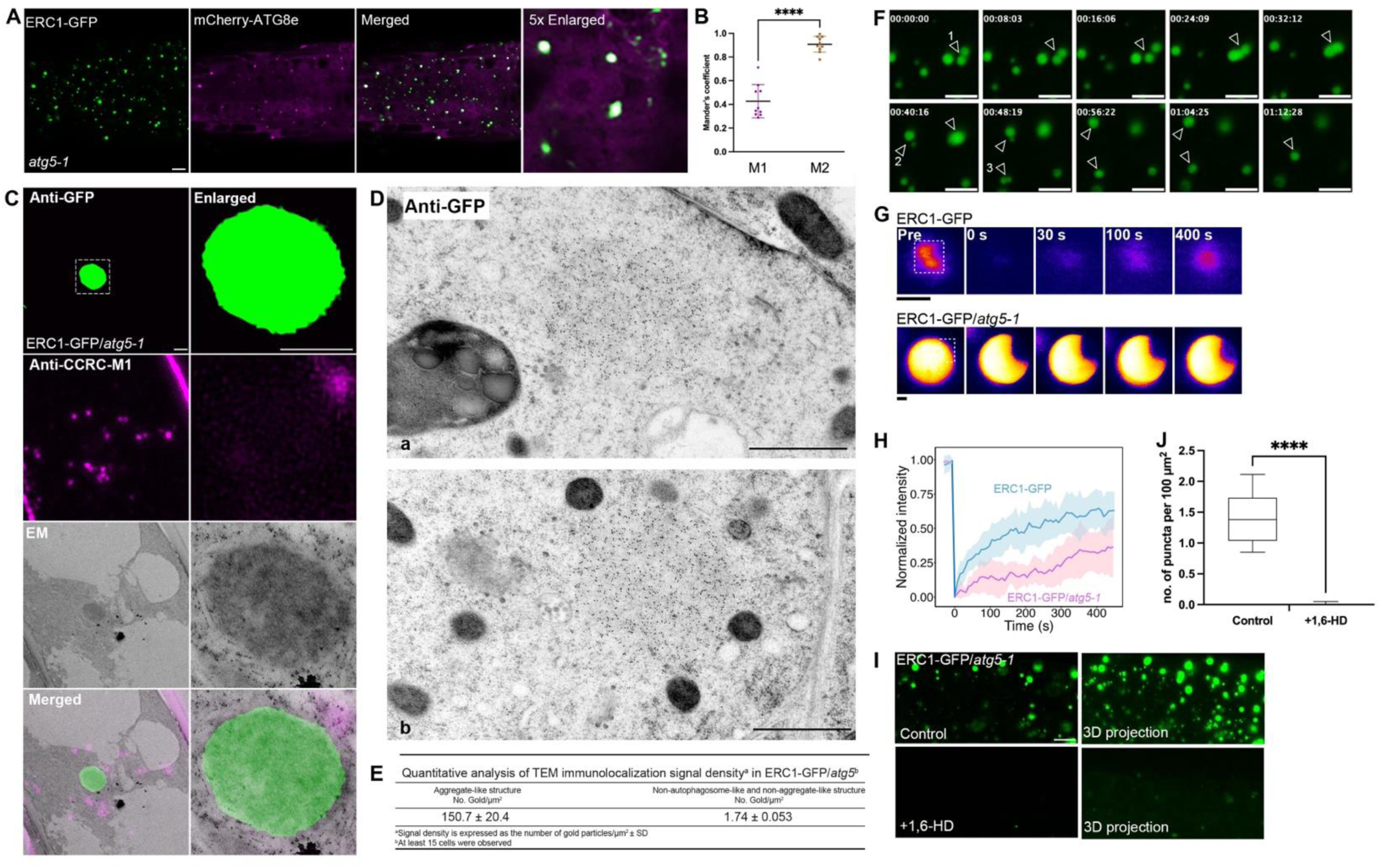
Accumulation of ERC1-GFP on large membraneless structures in core *atg* mutants. **(A)** Accumulation of mCherry-ATG8e on ERC1-GFP-positive large condensates in *atg5-1* mutant. Five-day-old seedlings of ERC1-GFP x mCherry-ATG8e were subjected to confocal imaging. Scale bar=10 μm. **(B)** Quantification of the Mander’s colocalization coefficients between ERC1-GFP and the mCherry-ATG8e from (A). M1, fraction of ERC1-GFP signal that overlaps with mCherry-ATG8e signal. M2, fraction of mCherry-ATG8e signal that overlaps with ERC1-GFP signal. Bars indicate the mean ± SD of 10 replicates. **(C)** Correlative light and electron microscopy (CLEM) analysis of ERC1-positive large puncta in Arabidopsis *atg5-1* mutant background. The left column showed the alignment of the confocal image (anti-GFP against ERC1-GFP in green and anti-CCRC-M1 in magenta against cell wall signal) with the TEM micrograph using the cell wall profile (anti-CCRC-M1). The right column showed an enlarged image for the cropped region indicated by the dash box. Scale bar=1 μm. **(D)** Immunolabeling with GFP antibodies showed accumulation of ERC1-GFP on large membraneless spherical structures in Arabidopsis *atg5-1* mutant. 5-days-old root tips of ERC1-GFP/*atg5-1* were subjected to high-pressure freezing fixation and sections were immunolabeled with anti-GFP antibodies. Scale bar=1 μm. **(E)** Quantitative analysis of immunogold labeling density in ERC1-GFP/*atg5-1* under TEM. **(F)** Time-lapse imaging illustrated fusion events of ERC1-positive condensates indicated by arrow and arrowhead in 5-day-old ERC1-GFP/*atg5-1* root cells. Scale bar=10 μm. (**G-H**) FRAP analysis of the ERC1-positive condensates in 5-day-old ERC1-GFP and ERC1-GFP/*atg5-1* root cells. Dashed box, bleached condensate. Quantification of the results was shown in (H). Data indicate the mean ± SD of 6 replicates. Scale bar=1 μm. (**I-J**) ERC1-positive condensates were disrupted by 1,6-hexanediol (1,6-HD). 5-day-old ERC1-GFP/*atg5-1* seedlings were treated without (control) or with 1,6-hexanediol (+1,6-HD). Quantification of the number of ERC1-GFP puncta was shown in (J). Scale bar=10 μm.

To access the morphological feature of these ERC1-positive condensates, we performed correlative light and electron microscopy (CLEM) analysis using the HPF fixed *atg5-1* transgenic plants expressing ERC1-GFP. As shown in **Figure 5C**, the GFP signal corresponding to ERC1-GFP was aligned to a dense membraneless structure under the EM. Consistently, immuno-EM analysis in ERC1-GFP/*atg5-1* root cells using gold particles against GFP antibodies also showed that gold particles were extensively decorated on large spherical sponge-like structures without a recognizable membrane boundary (**Figure 5D and 5E**).

The above observation promoted us to further access the property of the ERC1-positive condensates. Time-lapse imaging and fluorescence recovery after photobleaching (FRAP) are classical methods for studying the internal mobility of liquid droplets (Fang et al., 2019; Hatzianestis et al., 2023; Dragwidge et al., 2024; Miao et al., 2024; Mosesso et al., 2024). We observed coalescence of several ERC1-positive condensates in *atg5-1* mutant plants indeed occurred, which leads to formation a larger condensate as illustrated in the montage analysis **(Figure 5F, Supplementary Movie 3)**. In addition, when FRAP was applied to monitor the dynamics of ERC1 proteins in these condensates, the FRAP curves for the indicated ERC1-positive condensate showed ∼50% of ERC1-GFP signals gradually recovered over time in the wild type background, while ∼25% recovery of ERC1-GFP/*atg5-1* was detected, suggesting the molecular mobility is reduced **(Figure 5G and 5H)**. Application of 1,6-hexanediol (1,6-HD), an alcohol to dissolve phase-separated condensates (Alberti et al., 2019; Field et al., 2023), also greatly interfered the signal of the ERC1-positive condensates **(Figure 5I and 5J)**.

To gain more insights into the spatial organization of these membraneless compartments, we next applied 3D electron tomography (ET) analysis to collect a serial section using the HPF fixed ERC1-GFP/*atg5-1* root samples. Two representative examples were shown in **Figure 6, Supplementary Movie 4 and 5**. It is notable that ER membranes (coded in blue) were frequently detected in close juxtaposition with the condensate structure (coded in white). After 3D modeling, it is more evident that the condensate structure is likely encapsulated by the ER membranes. Remarkably, small vesicle-like structures (<100nm) with dense intensity (coded in purple) were discretely embedded within the lumen of the condensate structure. In addition, several Golgi stacks and PVCs/MVBs with clear boundaries were also observed nearby the condensate.

**Figure 6.**
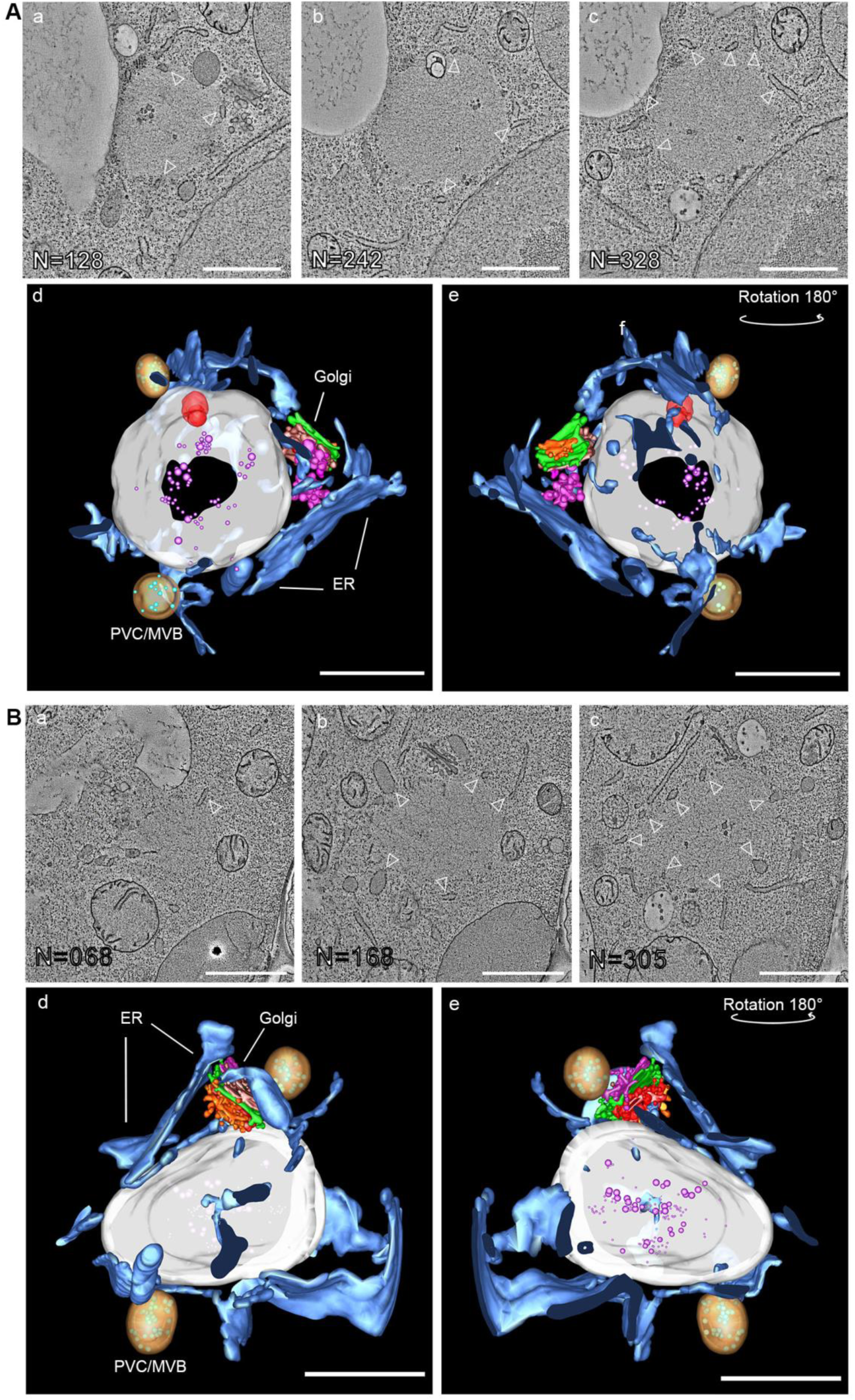
3D electron tomography analysis reveals the spatial relationship between the ERC1-postive membraneless structure and other membranes in *atg5-1*. (**A**, a-c) and (**B**, a-c) showed a gallery of the tomographic images displaying two examples of the large membraneless structures in ERC1-GFP/*atg5-1* root cells. Bottom panel in (A, d-e) and (B d-e) showed the models of 3D electron tomography depicting the spatial organization of the ERC1-postive membraneless structure (in white) and the endomembrane structures. The ER elements surrounding the membraneless structure were shown in blue. Small vesicles found within the ERC1-GFP membraneless structure were coded in purple. Scale bar=1 μm.

### ERC1 binds to NBR1 and responses to heat stress condition

Recent studies have suggested that biomolecular condensation of SARs including P62 and NBR1 occurs prior to phagophore initiation as an assembly platform for phagophore and ATG machinery (Agudo-Canalejo et al., 2021; Feng et al., 2023). Since ATG8e proteins were directed to ERC1-positive condensate structures in the absence of ATG8 conjugation activities (**Figure 5**), we wonder whether ERC1-positive condensates might function as a nucleation site for recruiting cargo or substrates prior to the occurrence of ATG8 conjugation onto the phagophore membrane. Hence, we generated Turbo-ID tagged ERC1/*atg7-2* transgenic plants and conducted a proximity labeling assay to search the potential molecules associated with the ERC1-positive condensates. As shown in **Figure 7A**, TurboID-coupled mass spectrometry analysis showed that in addition to ATG8e, another well-known SAR, NBR1, was significantly enriched. Interestingly, SH3P2, another ATG8-binding protein which contains a BIN-amphiphysin-RVS (BAR) domain essential form membrane deformation (Zhuang et al., 2013; Ahn et al., 2017; Sun et al., 2022), was also identified in the list. To access the possible relationship between ERC1 and NBR1, we firstly carried out a recruitment assay using CNX-mCherry-ERC1 and YFP-NBR1. Notably, YFP-NBR1 was redirected by CNX-mCherry-ERC1, but not by CNX-RFP (**Figure 7B**). Meanwhile, Co-IP analysis using HA-tagged NBR1 and ERC1-GFP also showed they were associated with each other (**Figure 7C**). We next sought to examine their possible direct interaction via a pull-down analysis. As shown in **Figure 7D**, His-tagged UBA domain of NBR1 is sufficient to bind to GST-ERC1.

**Figure 7.**
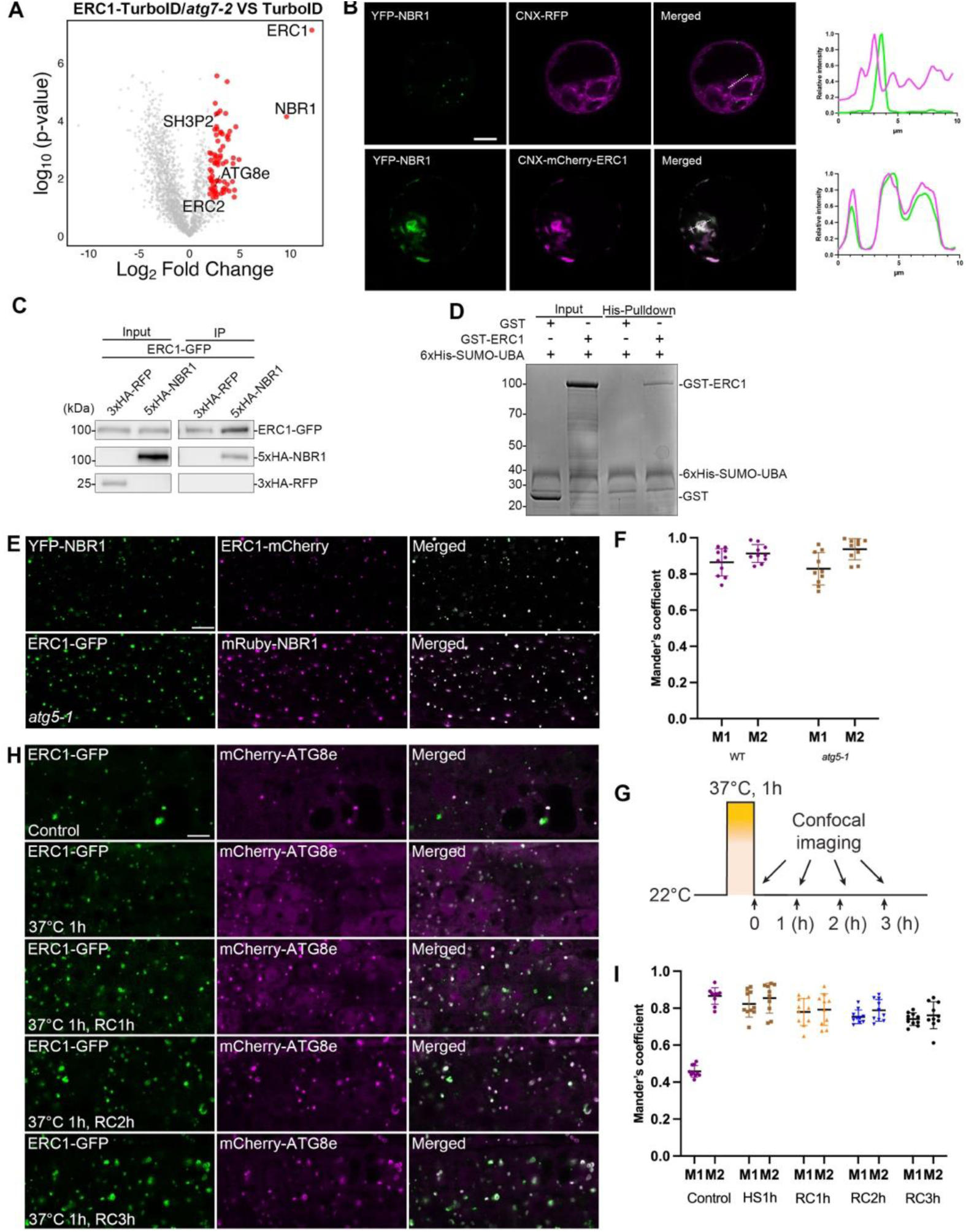
ERC1 interacts with NBR1 and responses to heat stress. **(A)** Volcano plots showing the proximity proteomics between ERC1-TurboID/*atg7-2* and TurboID control. Enriched autophagy-related proteins are highlighted with annotations. **(B)** Recruitment assay of YFP-NBR1 with CNX-RFP or CNX-mCherry-ERC1 in transient expressed Arabidopsis protoplasts. CNX-mCherry-ERC1 or CNX-RFP and YFP-NBR1 were transiently co-expressed in Arabidopsis protoplasts and subjected to confocal observation. Scale bar=10 μm. Similar results were obtained from three different independent experiments. Fluorescence intensity profiles were calculated along the white dash lines indicated in merged channels to show signal distribution of YFP-tagged NBR1 and CNX-RFP/CNX-mCherry-ERC1. **(C)** IP analysis showed that ERC1 is associated with with NBR1. Cell lysate from Arabidopsis PSBD protoplasts transiently expressing 3xHA-RFP or 5xHA-NBR1 with ERC1-GFP were subjected to GFP-trap assay. Input and IP samples were detected with anti-GFP and anti-HA antibodies respectively. Similar results were obtained from three different independent experiments. **(D)** His pull-down of 6xHis-SUMO-UBA with GST or GST-tagged ERC1 truncations. 10% of input and pulldown fraction was loaded. The input and pulldown fraction were visualized by Coomassie Blue staining. Similar results were obtained from three different independent experiments. **(E-F)** ERC1 colocalizes with NBR1 on large condensate structures in *atg5-1*. 5-day-old Arabidopsis seedlings expressing YFP-NBR1 x ERC1-mCherry or ERC1-GFP x mRuby-NBR1/*atg5-1* were subjected to confocal analysis. Scale bar = 10 mm; Quantification of the Mander’s colocalization coefficients between ERC1 and the NBR1 was shown in (F). M1, fraction of ERC1 signal that overlaps with NBR1 signal. M2, fraction of NBR1 signal that overlaps with ERC1 signal. Bars indicate the mean ± SD of 10 replicates. **(G-I)** HS treatment induced the formation of ERC1 puncta that colocalized with mCherry-ATG8e. 5-day-old Arabidopsis seedlings were incubated in preheated liquid ½ MS medium at 37 °C for 1 hour, followed by confocal microscope observation after recovery at the indicated time points shown in (**G**). Scale bar = 10 mm. Quantification of ERC1-GFP and mCherry-ATG8e colocalization was shown in (**I**). M1 is the fraction of the ERC1-GFP signal that overlaps with the mCherry-ATG8e signal; M2 is the fraction of the mCherry-ATG8e signal that overlaps with the ERC1-GFP signal.

To further investigate the dynamic relationship between NBR1 and ERC1, we generated transgenic plants expressing YFP-NBR1 x ERC1-mCherry in the wild type and *atg5-1* background respectively. Surprisingly, under the normal conditions, signals of YFP-NBR1 were well overlapped with that of the ERC1-positive puncta **(Figure 7E, top panel)**. On the other hand, in the absence of ATG5, hyperaccumulation of NBR1 on ERC1-positive large condensates was also observed **(Figure 7E, bottom panel)**. Several studies have reported that NBR1 actively participates in autophagy for heat resistance (Zhou et al., 2013; Zhou et al., 2014; Jung et al., 2020a; Thirumalaikumar et al., 2021), we thereby tested whether ERC1 responses to heat stress conditions. When 5-day-old ERC1-GFP x mCherry-ATG8e seedlings were subjected to mild heat stress under 37°C for 1h, we observed that numbers of ERC1-GFP puncta increased with various sizes and forms, and closely associated with mCherry-ATG8e on open or closed ring-like structures, as well as condensate-like structures during the recovery period **(Figure 7H-G, Supplementary Figure S4).**

### Deficiency of ERC1 compromises the turnover of the autophagic receptor NBR1 for stress resistant

To further understand the physiological role of ERC protein family in *Arabidopsis*, we screened the T-DNA insertion lines for ERC1 and ERC2 respectively (**Supplementary Figure S5**). RT-PCR analysis showed that transcripts were barely detected in *erc1-1, erc1-2 and erc2-1* mutants, but only a mild knockdown effect was observed in *erc2-2* mutant (**Supplementary Figure S5C**). We therefore used *erc1-1* and *erc2-1* for the following phenotype analysis. Under normal conditions, there was no distinguishable phenotype in the mutants. Of note, when the seedlings were subjected to nitrogen starvation, it seemed that *erc1-1* was more sensitive to nitrogen starvation than the wild type, ERC1 overexpression line and *erc2-1* (**Supplementary Figure S5D**). On the other hand, when a prolonged heat stress treatment (37°C for 72h) followed by a recovery period (22°C for 72h) was applied, we also observed overall seedling survival rate was severely suppressed in the *erc1-1* mutant when compared with that in the wild type, ERC1 overexpression line (ERC1-OE) and *erc2-1* (**Figure 8A and 8B**).

**Figure 8.**
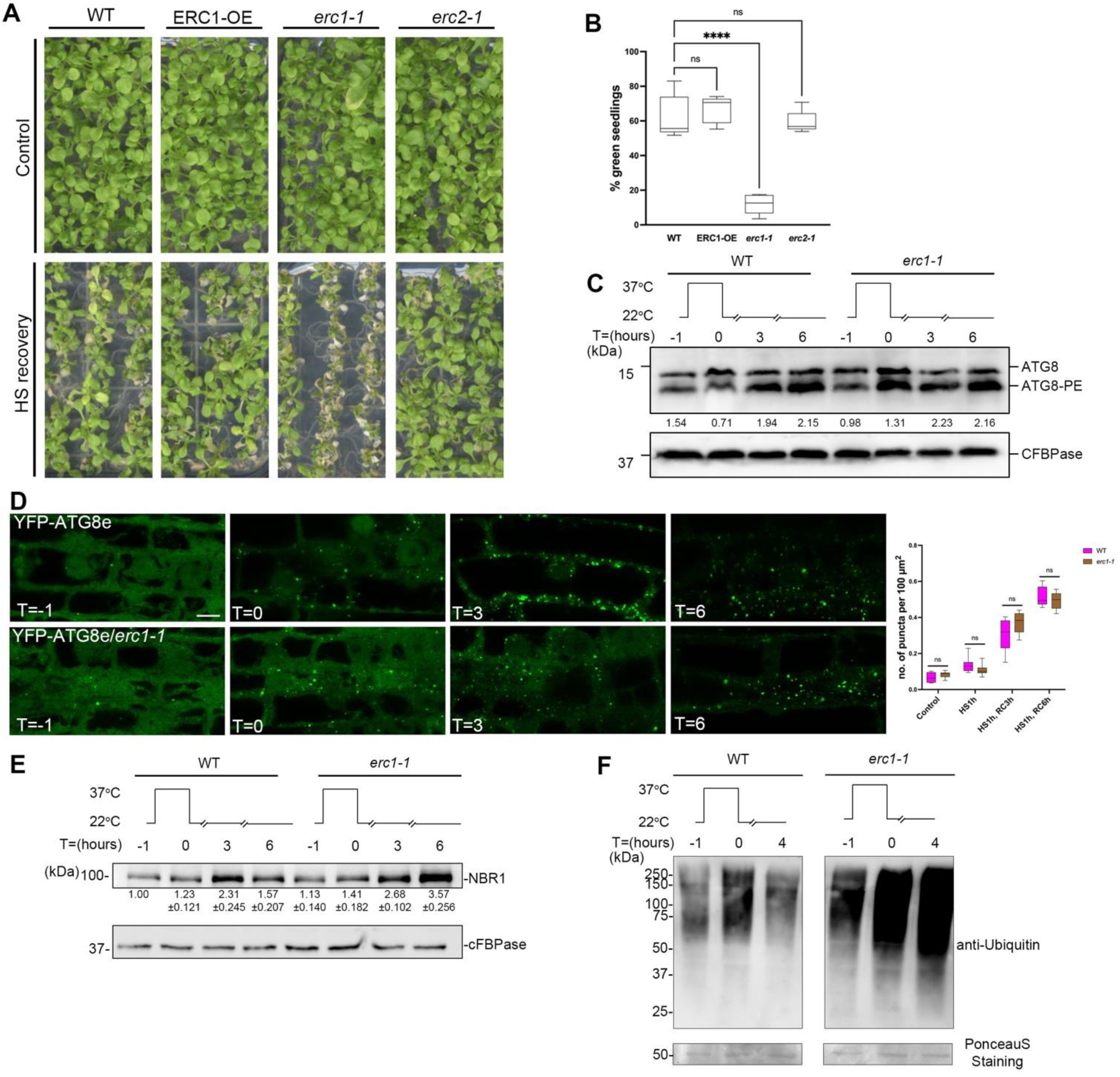
ERC1 deficiency compromises NBR1 turnover to improve heat recovery in Arabidopsis. **(A)** Phenotypic analysis under HS condition. 5-day-old Arabidopsis seedlings were subjected to 37 °C treatment for 3 days, then recovered at physiological growth temperature (22 °C) for 7 days. Similar results were obtained from three different independent experiments. **(B)** Data represents the percentage of green seedlings of Wild type (control), ERC1 overexpression line (ERC1-OE), *erc1-1*, and *erc2-1*. Five independent experiments were performed. Means ±SD with asterisks is significantly different at P < 0.0001 (one-way ANOVA). **(C)** ATG8 lipidation analysis of WT and *erc1-1* during heat shock treatment. 5-days-old Arabidopsis seedlings were incubated under 37 °C for 1 hour, followed by recovery under 22 °C for indicated periods. Membrane fractions were extracted from the indicated periods and subjected to immunoblot with anti-ATG8 antibodies. cFBPase was used as a loading control. The value represents the ATG8-PE vs. ATG8. **(D)** Confocal analysis of YFP-ATG8e in WT and *erc1-1* root cell upon heat stress. 5-day-old Arabidopsis seedlings were incubated in preheated liquid 1/2 MS medium under 37 °C for 1 hour, followed by confocal microscope observation after recovery under 22 °C for indicated periods. Scale bar = 10 μm. Quantification of the number of YFP-ATG8e puncta in wild type (WT) and *erc1-1* was shown in (**E**). Means ±SD are shown. **(E)** Immunoblot analysis of NBR1 turnover upon HS treatment. 5-day-old Arabidopsis seedlings were incubated in preheated liquid 1/2 MS medium under 37 °C for 1 hour and total proteins were extracted and immunoblotted with anti-NBR1 antibodies. cFBPase was used as a loading control. The value represents the relative protein content, defining the control group as 1. Data represent mean ± SEM. Similar results were obtained from three different independent experiments. **(F)** Ubiquitiantion level of insoluble proteins in WT and *erc1-1* upon HS treatment. Insoluble proteins were extracted from of 5-day-old WT and *erc1-1* plants from indicated time points upon HS treatment and subjected to immunoblot with anti-ubiquitin antibodies. PonceauS staining was used as a loading control. Similar results were obtained from three different independent experiments.

Autophagic flux can be monitored by ATG8 lipidation assay via measuring the cytosolic ATG8 and membrane-conjugated ATG8-PE form using immunoblot with ATG8 antibodies (Liu and Bassham, 2012; Marshall and Vierstra, 2018). Thereby, to further evaluate whether ERC1 regulate autophagic flux, we performed an immunoblot analysis to detect the ATG8 lipidation level before and after heat stress recovery. It appeared that ATG8-PE level was upregulated in both wild type and *erc1-1* backgrounds after HS (**Figure 8C**). To further assess whether the number or shape of the autophagosomal structures is disturbed, we introduced YFP-ATG8e into *erc1-1* mutant for confocal analysis. Consistently, number of YFP-ATG8e foci increased in both WT and *erc1-1* after HS recovery (**Figure 8D and 8E**). After 6 hours recovery, it appeared that autophagosome structures labeled by YFP-ATG8e were slightly increased in the wild type than in *erc1-1* mutant.

As ERC1 also interacts with NBR1, which has been reported to target ubiquitinated protein aggregates for clearance via autophagy under heat stress (Zhou et al., 2013; Zhou et al., 2014; Jung et al., 2020a; Thirumalaikumar et al., 2021), we sought to examine the impact of ERC1 deficiency for the turnover of NBR1 and total ubiquitination level during HS recovery (**Figure 8F and 8G**). As shown in **Figure 8F**, when loss of ERC1, NBR1 level was upregulated in *erc1-1* mutant after 3h HS recovery (**Figure 8F**), suggesting that ERC1 is required for NBR1-mediated autophagic degradation during HS recovery. In agreement with this result, immunoblot analysis showed that ubiquitinated proteins were more abundant in *erc1-1* than in wild type after HS recovery (**Figure 8G)**.

## Discussion

Autophagy requires a sophisticated regulatory network to exert accurate response to different environmental stimuli for maintaining cellular homeostasis. In this study, we have identified *Arabidopsis* ERC1 as a novel plant autophagic regulator to promote autophagosome formation via biomolecular condensation. We demonstrated a physical interaction between *Arabidopsis* ERC protein family and ATG8e in an AIM-dependent manner **(Figure 1 and 2)**. Interaction mapping in combination with a complex modeling uncovered an “WDFF” motif in the extended C-terminus of ERC1 is essential for ATG8 interaction **(Figure 2).** The first “W” is a canonical residue conserved in many canonical AIM sequences (Kirkin and Rogov, 2019). Interestingly, the fourth “F” has only been found in the AIM sequence of ATG30, a SAR for pexophagy in yeast *Pichia pastori* and its other homologs, and its interaction with ATG8 is phosphoregulatable (Farre et al., 2008; Farre et al., 2013). Therefore, the “WDFF” of ERC1 might represent a diversified AIM to confer ATG8-binding specificity in plants. On the other hand, we still observed that signals of ERC1 condensates were encapsuled by CNX-RFP-ATG8e when the “WDFF” was removed or substituted, but an ERC1 truncation lacking a predicted IDR region (T3) changed to a punctate pattern which was restricted to the peripheral region of CNX-RFP-ATG8e (**Figure 2B**). One possible explanation is that phase separation of ERC1 also attribute to its binding strength with ATG8, while highly avid interactions would be achieved via ERC1 condensation to promote autophagic degradation.

We further demonstrated that independent of the ATG8 conjugation system, ERC1 is capable to from liquid-like condensates with ATG8 and NBR1 proteins (**Figure 5 and Figure 7**). Previous studies have indicated that ATG8-positive large foci were occasionally observed in *atg4a/b* and *atg7* mutants, which are suggested to be protein aggregates (Yoshimoto et al., 2004; Jung et al., 2020a). Our results revealed ERC1 condensates exhibit liquid-like behavior including fusion and diffusion (**Figure 5F-I**). In addition, ATG8 and NBR1 proteins are uniformly distributed in the ERC1 condensates, rather than associated only with the outer layer as a ring-like pattern, supporting the fluidity within the ERC1 condensates (**Figure 5A and Supplementary Figure S3**). Furthermore, our ultrastructural dissection unveils a distinct spatial architecture of the ERC1-dependent condensates with other membrane structures. Morphologically, ERC1 condensate structures are membraneless, but closely associated with ER membranes (**Figure 5D and Figure 6, and Supplementary Movie 3 and 4**). Remarkably, many vesicles are clustered within the ERC1 condensates, highlighting a multiphasic architecture of ERC1 condensates that has not been described in plant cells. It is tempting to speculate that prior to ATG8 conjugation, ERC1 would promote ATG8 nucleation via phase separation to facilitate membrane nucleation (e.g. ER and vesicles), and subsequent membrane remodeling. The interaction between ERC1 and the autophagic receptor NBR1 further outlines a possible mechanistic framework for cargo sequestration via ERC1 and NBR1 condensation before phagophore initiation (**Figure 7**). In mammalian cells, protein condensates have been reported to interact with membranes via a “wetting” process to recruit membrane sources and induce membrane remodeling for autophagosome formation in animal cells (Agudo-Canalejo et al., 2021; Feng et al., 2023; Mangiarotti and Dimova, 2024). Several latest findings have also highlighted a close interplay between biomolecular condensation and membranes in plants. It was reported that a component of processing bodies, DCP1 is associated with plasma membrane to modulate the condensation of SCAR-WAVE complex for actin nucleation (Liu et al., 2023). A recent study has demonstrated that phase separation by the endocytic TPLATE complex on the membrane promotes the recruitment of other endocytic adaptors for clathrin-coated vesicle formation (Dragwidge et al., 2024). An endosomal sorting complex required for transport (ESCRT) protein, ALIX undergoes phase separation together with CaLB1 to facilitate autophagosome closure (Mosesso et al., 2024). Additionally, phase separation of VPS41, a homotypic fusion and protein sorting (HOPS) subunit, has also been implicated to induce the formation of VPS41-associated phagic vacuoles to modulate autophagy degradation (Jiang et al., 2024). In future, it will certainly be worthwhile to further investigate the possible role(s) of ERC1 condensates in membrane source gathering or remodeling process for phagophore initiation. Undoubtedly, high-resolution and multidimensional assessment of biomolecular condensates will allow us to gain deeper insights into the interplay between membraneless condensates and membranes for plant stress adaptation.

Functionally, we found that suppression of ERC1 compromises plant resistance to heat stress and nutrient deprivation (**Figure 8A and Supplementary Figure S5D**). It seemed that the ATG8 lipidation level and number of autophagosomal structures were comparable in the WT and *erc1-1* mutant (**Figure 8C-D**). A recent study also reported that autophagic flux in a *nbr1* mutant is indistinguishable from the WT upon heat stress (Jung et al., 2020a). Our data showed that found that dysfunction of ERC1 leads to accumulation of NBR1 and the total ubiquitinated proteins (**Figure 8F and 8G**). It is very likely that ERC1 functions in NBR1-dependent autophagic degradation but is dispensable for bulk autophagy as NBR1. Nevertheless, we can’t exclude the possibility that other alternative ERC-independent autophagic pathway (s) is induced under heat stress when loss of ERC1 activity. Significant progress has been made to underpin the vital roles of membraneless condensates as thermosensors for plant heat stress resistance (Jung et al., 2020b; Zhu et al., 2022; Blagojevic et al., 2024), but whether and how they are coupled with membrane dynamics is unknown. It is surprising that the ERC protein family shares high homology in the coil-coil domains with its orthologs in most representative land plant species, while its functions in other plants have not been described. More efforts are required in future to elucidate the interaction network of ERC protein family in plants. Collectively, this work offers a ERC1-medidated biomolecular condensation model for ATG8 and NBR1 recruitment in response to plant heat stress, providing new insight into the assembly mechanism for plant autophagosome initiation.

## Materials and Methods

### Plasmid construction

The GFP/YFP/RFP/mCherry/Flag-tagged constructs used for *Arabidopsis thaliana* Col-0 transgenic plants or transient expression in protoplasts were built by cloning through the insertion of PCR-amplified cDNA into the pBI121/221/pCAMBIA1300 backbone driven by the UBQ10 promoter. To generate CNX-mCherry-ERC1, a ERC1 fragment with stop codon and a NOS fragment was fused by fusion PCR, followed by insertion into 221-CNX-mCherry-ATG8f (Sun et al., 2022) digested by SpeI/EcoRI. 3Flag-3HA-SH3P2-NOS fragment was obtained by fusion PCR and subcloned into pBI221/pBI121 to generate pBI221/pBI121-3Flag-3HA-SH3P2, followed by insertion with TurboID-ATG8 digested by XbaI/EcoRI, or TurboID digested by XbaI/SalI. The 3Flag-3HA-TurboID fragment was digested by BamHI/EcoRI and inserted into pCAMBIA1300-YFP-ATG8f to generate pCAMBIA-3Flag-3HA-TurboID. For 1300-UBQ-ERC2-TurboID, a TurboID-HA-NOS fragment was obtained by fusion PCR and inserted into pCAMBIA-1300-ERC1-GFP digested by KpnI/EcoRI. A synthesized mRuby fragment was inserted into the pCAMBIA1300-YFP-ATG8f (Zhuang et al., 2013) digested by BamHI/SpeI to generate a pCAMBIA1300-mRuby-ATG8f. The coding sequence of NBR1 was then subcloned into the digested pCAMBIA1300-mRuby-ATG8f by SpeI/KpnI to generate pCAMBIA1300-mRuby-NBR1. For *E.coli* expression constructs, the full-length/truncations coding sequence of ERC1 or ATG8e were cloned into pGEX-6P-1 or pET30a+ respectively via restriction digestion and ligation to generate GST-tagged ERC1/truncations, and 6xHis-ATG8e. GST-ATG8e construct was described previously (Zhuang et al., 2013). 221-YFP-ATG8e, 221-YFP-ATG8f, 221-YFP-NBR1 and 221-CNX-RFP were described previously (Sun et al., 2022). All constructs were verified by sequencing. Constructs and primers used in this study are listed in Supplementary Table 1.

### Plant materials

In this paper, all plant materials were the *Arabidopsis thaliana* Col-0 background. *Arabidopsis thaliana* T-DNA insertional mutants *erc1-1* (SAIL_129_D04)*, erc1-2* (SALK_112206), *erc2-1* (GK_151G05), *erc2-2* (SALK_059135) were obtained from the Nottingham Arabidopsis Stock Centre. The marker lines YFP-ATG8e, mCherry-ATG8e, ER-mCherry, ST-RFP1, mCherry-Rha1, VHA-a1-RFP1, YFP-NBR1, and mutants *atg2-1* (SALK_076727), *atg5-1* (SAIL_129_B07) *and atg7-2* (GK-655B06), have been described previously (Zhuang et al., 2017; Luo et al., 2023). ERC1-GFP, ERC1-mCherry, mRuby-NBR1, ERC1-GFP/*atg2-1,* ERC1-GFP/*atg5-1,* ERC1-GFP*/atg7-2,* ERC1-GFP×mCherry-ATG8e*/atg5-1* and YFP-ATG8e*/erc1-1,* were generated by floral dip method (Clough and Bent, 1998). ERC1-GFP× mCherry-ATG8e, ERC1-GFP×ST-RFP1, ERC1-GFP×VHA-a1-RFP1, ERC1-GFP×mCherry-Rha1, ERC1-GFP×mRuby-NBR1/*atg5-1,* were generated by crossing. All the transgenic lines were screened using antibiotic selection, followed by fluorescence signal checking under the microscope.

### Plant growth and treatment conditions

*Arabidopsis* seeds were surface-sterilized and then sown on the plates containing ½ Murashige and Skoog (MS) salts, 1% (w/v) sucrose, and 0.8% (w/v) agar (pH 5.7). The plates were placed at 4 °C for 48-72 h and then moved to the growth chamber at 22 °C under a long-day (16 h light and 8 h dark) photoperiod. Plants exposed to long-day conditions were transferred to soil after 1 week. For chemical treatments, 4- or 5-day-old seedlings were treated with liquid ½ MS medium as a control, or liquid ½ MS medium containing 100 μM BTH (Sigma), 10 μg/mL BFA (Sigma), 8.95 μM wortmannin (Sigma), or 0.5 mM Conc A (Santa Cruz) for the indicated periods before observation. For nitrogen starvation, around 50 sterilized and stratified seeds were transferred to liquid ½ MS medium in 6-well plates and placed on a shaker (60-80 rpm) for 7 days under continuous lighting. After 7 days, seedlings were washed with sterilized water three times and replenished with liquid ½ MS medium (control) or liquid ½ nitrogen-free MS medium (Caisson). The seedlings were put back into the growth chamber and incubated with shaking for an extra 14 days before observation. For heat shock experiments, 5-day-old seedlings were incubated in preheated liquid ½ MS medium at 37 °C for 1h, and then transferred to fresh ½ MS plates under 22 °C with the indicated periods for recovery. For the 1,6-Hexanediol assay, seedlings were submerged into liquid ½ MS medium with 15 % (w/v) 1,6-hexanediol and subjected to vacuum suction for 5 minutes, followed by 5 minutes of incubation before microscope observation.

### Transient expression in protoplasts

Methods for transient expression in *Arabidopsis thaliana* plant system biology dark type culture (PSBD) were performed as previously described (Miao and Jiang, 2007). In brief, 5-day-old PSBD cells were digested for 2 h and centrifuged at 100 × *g* for 10 min, followed by washing twice before electroporation. After transformation, protoplasts were incubated for 12 to 16 h in the dark before confocal imaging or protein extraction.

### Phylogenetic and sequence motif analysis

Sequences of the ERC protein family from 13 representative plant species were downloaded from the PANTHER gene ontology database (http://pantherdb.org/index.jsp). Protein sequence alignment was performed using Muscle v.3.8.31(Edgar, 2004). Phylogenetic trees were reconstructed by maximum likelihood using MEGA7(Kumar et al., 2016) with the 1000 bootstrap replicates and Evolview v3 (Subramanian et al., 2019). Conserved motif analysis within the representative ERC orthologs was performed via an online tool MEME suite (https://meme-suite.org/meme/index.html) as described (Bailey et al., 2009).

### Confocal imaging

Leica Stelleris8 confocal microscopes were used for detecting various fluorescence-tagged proteins with a HC PL APO 63×/1.2NA CORR CS2 water lens. GFP, YFP, and mCherry/RFP/mRuby were excited with 488/514/561 nm lasers, respectively. For each experiment or construct, more than 30 individual cells or individual 10 plants were observed for confocal imaging that represented >75% of the samples showing similar expression levels and patterns. For collecting 3D projection images, the depth (Δ*Z*) was set to 0.6 µm with total 18 stacks. Images and videos were further analyzed using LAS X, Adobe Photoshop software, and Fiji.

### Fluorescence recovery after photobleaching (FRAP)

Fluorescence recovery after photobleaching (FRAP) was performed using a Leica SP8 confocal laser scanning microscope equipped with a 488 nm laser. For FRAP experiments, six independent experiments were conducted and averaged to obtain a single curve. 5-days-old seedlings were transferred to an imaging chamber (Grace Bio-Labs) along with 1% (w/v) low-melting agarose (ThermoFisher). The imaging chamber was sealed by transparent sealing membrane to improve imaging stability. The FRAP parameters were as follow: 4 pre-bleaching images were recorded followed by 5 iterations of 488 nm set at 100% transmission. Pre-bleaching and recovery images (512 pixels × 512 pixels) were recorded for every 7.5 s for a total of around 400 s after bleaching. At each time point, maximum intensity projections from a z-stack of 10 steps with a step size of 1-2 μm were applied. Analysis of the recovery curves was carried out with Fiji and RStudio (Version 2024.04.1+748). The normalized fluorescence intensity was calculated as previously described (Zhou et al., 2023) with the following equation: *I_norm_ = [(I_recovery_ − I_bleach_)/(I_pre-bleach_ − I_bleach_)] × 100,* where *I_norm_* is the normalized intensity, *I_recovery_* is the recovery intensity after photobleaching, *I_bleach_* is the photobleaching intensity and *I_pre-bleach_* is the intensity before photobleaching.

### Spinning disk confocal microscopy

Spinning disk system used a Yokogawa CSU-W1 scanning unit with an Axio Observer Inverted microscope (Zeiss, 3i) and an ORCA-Fusion BT Digital C15440 sCMOS camera (Hamamatsu Photonics). A 63X objective (Plan-Apochromat 63X/1.15 Water Corr M27, Zeiss) was used. SlideBook 6 X64 software (version 6.0.17) was used to record time-lapse imaging. The parameters for spinning disk dynamic analysis were as follows: c488 and c560 lasers were used to excite green (GFP) and red (mCherry) channels respectively. 3D time-lapse were recorded with laser intensity 60-80% and exposure time 100-150 ms depending on the samples. The total 3D (z-stack) volumes were set to 15-20 μm with a step size 0.5 μm. The 4D movies were first processed to compensate photobleaching. Afterwards, deconvolution was perform followed by maximum intensity projection along the z-axis. All movie analysis were performed in SlideBook with default settings. The montages of the movies were made in ImageJ/Fiji.

### Protein extraction and immunoblotting

For total protein detection and ATG8 lipidation assay, 4- or 5-day-old seedlings were transferred to ½ MS liquid medium as control, or HS treatment with indicated recovery periods, and homogenized thoroughly in liquid nitrogen. The crude lysates were boiled in lysis buffer containing 100mM Tris-HCl, pH7.4, 20%(v/v) glycerol, 4%(w/v) SDS, 1%(v/v) beta-mercaptoethanol, and 0.8%(w/v) bromophenol blue for 10 minutes, followed by centrifugation at 15000 x g for 10 mins to remove cell debris. The resulting supernatants were separated by 10% SDS-PAGE gel and immunoblotted with anti-cFBPase (Agrisera), anti-GFP antibody (ABclonal) or anti-NBR1(Agrisera). For ATG8 lipidation assay, 4- or 5-day-old seedlings were subjected to HS treatment with indicated recovery periods. Seedlings were homogenized and boiled in lysis buffer containing 100mM Tris-HCl, pH7.4, 20%(v/v) glycerol, 4%(w/v) SDS, 1%(v/v) beta-mercaptoethanol, and 0.8%(w/v) bromophenol blue for 5 minutes, followed by centrifugation at 15000 x g for 10 mins to remove cell debris. The resulting supernatants were separated by 15% SDS-PAGE gel and immunoblotted with anti-cFBPase (Agrisera), or anti-ATG8 (Agrisera). For insoluble ubiquitinated protein assay, 4 or 5 days old seedlings were subjected to heat stress and recovery, followed by protein extraction using lysis buffer (50 mM Tris-HCl, pH 7.4, 150 mM NaCl, and 1mM EDTA) containing 1× Complete Protease Inhibitor Cocktail. The samples were collected and centrifuged at 1,000 × g for 5 minutes at 4 °C. The supernatant was then transferred to a new 1.5ml Eppendorf tube and centrifuged at 20,000 × g for 30 minutes. The insoluble fractions were re-suspended in 1x protein loading buffer. Protein samples were separated by 10% SDS–PAGE followed with immunoblotting analysis using anti-Ubiquitin (Chemcruz) and Ponceaus Staining.

### Co-Immunoprecipitation

*Arabidopsis* protoplasts were diluted 3-fold with 250 mM NaCl buffer before centrifuging at 100 × g for 10 min. Total cells were resuspended in 2× lysis buffer containing 25 mM Tris-HCl at pH 7.4, 150 mM NaCl, 1 mM EDTA, 0.4% (v/v) Nonidet P-40, and 1× Complete Protease Inhibitor Cocktail (Roche). The cells were homogenized using a 1 ml syringe with needle, followed by centrifugation at 13,000 × g for 20 min at 4 °C. The supernatant was collected and incubated with GFP-Trap beads (ChromoTek) with rotation at 4 °C for 3 h. Samples were washed in the wash buffer containing 25 mM Tris-HCl at pH 7.4, 150 mM NaCl, 1 mM EDTA, 0.4% (v/v) Nonidet P-40, for 4 times and then eluted by boiling in 1× protein loading dye. The samples were subjected to SDS-PAGE and immunoblotting with anti-Flag (Sigma) anti-HA (Sigma) and anti-GFP (ABclonal) antibodies.

### Electron tomography (ET), 3D reconstruction, and modeling

ET was performed using previously described methods (Zhuang et al., 2017; Ma et al., 2021; Luo et al., 2023). Briefly, 250-nm-thick sections were cut followed by poststaining using uranyl acetate and lead citrate. Sections were recorded using a Tecnai F20 electron microscope (FEI Company). For each grid, a tilt image stack from +60° to −60° with 1.5° increments was collected, and then the second tilt image stack was captured by rotating the grid by 90°. Dual-axis tomograms were calculated from pairs of image stacks with the etomo program of the IMOD software package (version 4.11). For 3D model generation, the contours were drawn manually and meshed with the 3dmod program in the IMOD software package.

### Correlative light and electron microscopy (CLEM)

ET was performed using previously described methods (Zhuang et al., 2017; Ma et al., 2021; Luo et al., 2023). Briefly, 250-nm-thick sections were cut followed by poststaining using uranyl acetate and lead citrate. Sections were recorded using a Tecnai F20 electron microscope (FEI Company). For each grid, a tilt image stack from +60° to −60° with 1.5° increments was collected, and then the second tilt image stack was captured by rotating the grid by 90°. Dual-axis tomograms were calculated from pairs of image stacks with the etomo program of the IMOD software package (version 4.11). For 3D model generation, the contours were drawn manually and meshed with the 3dmod program in the IMOD software package.

### CLEM

The root tips of 5-days-old seedlings were dissected and high-pressure frozen, followed by freeze substitution, HM20 embedding, and UV polymerization as described above. 150 nm sections were prepared and collected using formvar-coated nickel slot grids. The collected grids were incubated with anti-GFP and CCRC-M1 antibodies (Freshour et al., 2003). After washing, the grids were incubated in secondary antibody conjugated with Alexa Fluor 488 (ERC1-GFP signal) and 568 (cell wall profile). Afterward, the grids were visualized under a Leica Stellaris 8 confocal laser scanning microscope as described above. Subsequent TEM imaging was performed using a Hitachi H-7650 transmission electron microscope. The confocal and TEM micrographs were aligned by using cell wall profiles.

### Quantification and statistical analysis

For puncta quantification, Fiji was used to calculate the number of puncta and size of puncta in each channel using the Analyze Particles function in Fiji. Mander’s colocalization analysis was performed using the methods described previously (Luo et al., 2023), and JACoP plugin (Bolte and Cordelières 2006) was used in Fiji for Mander’s coefficient M1 and M2 analyses. For montage analysis and the intensity measurement of the time-lapse images, the region was selected using a rectangular selection tool in Fiji, and the mean grey value is obtained across the time points as indicated. For CNX recruitment assay quantification, a 10µm line was drawn across the region of interest and profile plot function in Fiji was used to measure the signal intensity in different channels. The resulted graph was generated using Prism9. For western blot quantification, the functions in Gels under Analyze module were used in Fiji. Experiments were performed for at least 3 biological replicates. Data are presented as mean values ± SD or ± SEM. Student’s *t* test or one-way analysis of variance (ANOVA) was applied to analyze the significant differences between different columns. Differences with *P* < 0.05 were considered significant. The significance levels are as follows: \**P* < 0.05, \*\**P* < 0.01, \*\*\**P* < 0.001, and \*\*\*\**P* < 0.0001 and ns represents no significance. Prism 9 was used to analyze and plot the differences among samples. Letters above the bars indicated significant differences.

### Expression and purification of recombinant proteins in *E.Coli*

pET-30a-ATG8e, pGEX-6P, pGEX-6P-ATG8e, pGEX-6P-ERC1, pGEX-6P-ERC1T1, pGEX-6P-ERC1T2, pGEX-6P-ERC1T3, pGEX-6P-ERC1T6, pGEX-6P-ERC1T10 were transformed into *E.coli* BL21(DE3) and induced at OD600 1.0 with 0.4mM IPTG at 16°C for 16 hours for protein expression. To purify 6xHis-ATG8e fusion proteins, cells were lysed by sonication in lysis buffer (50mM imidazole in 1×PBS pH7.4). The lysates were subjected to centrifugation at 15000×g, 4°C for 45 minutes. The supernatant was collected as the soluble fraction. 6xHis-ATG8e proteins were purified using High-Affinity Ni-Charged resins (Genescript) and eluted with elution buffer (500mM imidazole in 1×PBS pH7.4). The elution was subsequently dialysed against 1×PBS overnight. Proteins were concentrated to 4mg/ml, and stored at −80°C. To purify GST, GST-ATG8e and GST-ERC1/truncations proteins, the cells were collected and resuspended in lysis buffer (5mM DTT in 1xPBS) for sonication. The lysates were subjected to centrifugation at 15000xg, 4°C for 45minutes. The supernatant was collected as the soluble fraction. The soluble fractions were incubated with glutathione resins (GenScript) for 16 hours at 4°C. The proteins were eluted with elution buffer (20mM reduced glutathione, 50mM Tris-HCl pH8.0) at room temperature and subjected to dialysis against 1xPBS for 16 hours at 4°C. Proteins were concentrated to 4 mg/ml, and stored at −80°C. To remove the GST tag, the proteins were incubated with home-made Precision Protease at 4°C for 16 hours in 1xPBS with 5mM DTT. The resulting mixture was subsequently incubated with gutathione resins (GenScript) for 2 hours at room temperature and followed by High-Affinity Ni-Charged resins to remove the protease. The flowthrough was collected and concentrated to around 4 mg/mL and stored at −80°C.

pHisSUMO-NBR1-UBA was transformed into *E.coli* C41 and induced at OD600 0.2 with 0.4mM IPTG at 25°C for 16 hours for protein expression and purified as previously described (Sun et al., 2022). Cells were lysed by sonication in lysis buffer (50mM imidazole in 1xTBS pH7.4). The lysates were subjected to centrifugation at 15000 x g, 4°C for 45 minutes. The supernatant was collected as the soluble fraction. HisSUMO-NBR1-UBA proteins were purified using High-Affinity Ni-Charged resins (Genescript) and eluted with elution buffer (500mM imidazole in 1xTBS pH7.4). The elution was subsequently dialysed against 1x PBS overnight. Proteins were concentrated to 4mg/ml, and stored at −80°C.

### *In-vitro* pull-down assay

For GST pull-down assay, 0.2 uM of GST-ATG8e or GST together with ERC1 were incubated with glutathione magnetic resin (GenScript) in the reaction buffer (1% NP40, 1xPBS pH7.4) at room temperature on a rotary platform for 1 hour. Input samples were taken after the incubation. The magnetic resins were washed with reaction buffer for 3 times and eluted by boiling the magnetic resin in 1x protein loading dye. The sample were separated using 10% SDS-PAGE gel and visualized by Coomassie Brilliant Blue Staining. For His pull-down assay, 0.2 uM of 6xHis-ATG8e and GST-ERC1 FL/T1/T2/T3/T4/2A were incubated with Ni-Charged NTA magnetic resins (GenScript) in the reaction buffer (50mM imidazole, 1% NP40, 1xPBS pH7.4) at room temperature on a rotary platform for 1 hour. Input samples were taken after the incubation. The magnetic resins were washed with reaction buffer for 3 times and eluted by boiling the magnetic resins in 1x protein loading dye. The sample were separated using 10% SDS-PAGE gel and visualized by Coomassie Brilliant Blue staining.

For His-SUMO-UBA pull-down assay, 0.2 uM of 6xHis-SUMO-UBA and GST/GST-ERC1 FL were incubated with Ni-Charged NTA magnetic resins (GenScript) in the reaction buffer (50mM imidazole, 1% NP40, 1xPBS pH7.4) at room temperature on a rotary platform for 1 hour. Input samples were taken after the incubation. The magnetic resins were washed with reaction buffer for 3 times and eluted by boiling the magnetic resins in 1x protein loading dye. The sample were separated using 10% SDS-PAGE gel and visualized by Coomassie Brilliant Blue staining.

### RNA Extraction and RT-PCR

RNA from 5-day-old seedlings were extracted according to manufacturer’s protocol (Purelink RNA Mini Kit, Invitrogen). The RNA was transcribed into cDNA using M-MLV reverse transcriptase kit (Promega) according to manufacturer’s protocol. The cDNA was subjected to PCR amplification with indicated cycles and separated with agarose gel electrophoresis. The primers were listed in Supplementary Table 1.

### Model building for ERC1 and ATG8e

The complex structure prediction of ERC1 peptide and ATG8e was generated by AlphaFold2 Multimer (Richard Evans, 2022) and implemented in ColabFold (Mirdita et al., 2022). Structural analysis and figure preparation were performed using UCSF ChimeraX (version 1.8.dev202404160016) (Pettersen et al., 2021).

### Proximity labeling coupled with mass spectrometry analysis

Procedures of biotin treatment and affinity purification were performed according to previously described methods (Mair et al., 2019) with minor modifications. Briefly, 1 g of 5-days-old seedlings expressing TurboID-ATG8/*atg2-1,* or ERC1-TurboID/*atg7-2,* and TurboID as control, were treated with 50 µM biotin for 1 h, respectively. The seedlings were quickly rinsed with water and then frozen in liquid nitrogen before affinity purification. For the affinity purification, 1 g of grounded plant powder was resuspended in 2 ml extraction buffer (50 mM TrisHCl pH 7.5, 150 mM NaCl, 0.1% SDS, 1% Triton-X-100, 0.5% Na-deoxycholate, 1 mM EGTA, 1 mM DTT, 1x complete Protease inhibition cocktail, 1 mM PMSF) followed by 15,000 g centrifugation at 4 °C for 15 mins. The clear supernatant was subjected to a 7k Zeba Spin desalting column (Thermo Fisher) to remove excess biotin, and then transferred into a new 5 ml tube containing 150 µl pre-washed bead slurry of Dynabeads MyOne Streptavidin C1 (Invitrogen), and supplemented with a final concentration of 1x complete protease inhibitor and 1 mM PMSF for incubation on a rotor wheel at 4°C for 16 h. The beads were then separated from the protein extract on a magnetic rack and washed as previous described and stored at −80°C (Mair et al., 2019). The MS sample preparation, LC-MS/MS, and data analysis were performed according to (Mair et al., 2019). The frozen beads were thawed for on-beads digestion, followed by peptide desalting using C18 desalting tips and dried in a speed vac, and stored at −80°C before LC-MS/MS. For quantitative comparisons, protein intensity values in three biological replicates were log_2_-transformed before further analysis. Proteins with less than two valid values in at least one group were removed and the missing values were replaced from a normal distribution with a width of 0.3 and a downshift value of 1.8 (the default values). Student’s t-test was conducted with a threshold of FDR = 0.05 and S0 = 0.5.

## Acknowledgments

This work was supported by grants from the National Natural Science Foundation of China (32222087), the Research Grants Council of Hong Kong (N_CUHK405/20, 24108820, 14106622, 14110923, G-CUHK408/23, C4002-20W, C4002-21EF, C4033-19E, R4005-18, AoE/M-05/12, and AoE/M-403/16), and The Chinese University of Hong Kong (CUHK) Research Committee to X.Z., and grants from the National Natural Science Foundation of China (32270291, 32061160467) to C.G, and the National Institutes of Health grant R01GM135706 and the Carnegie Endowment Fund to the Carnegie Mass Spectrometry Facility to S.L.X..

## Notes

### Competing Interest Statement

The authors have declared no competing interest.

