## Supplemental Figure 1-5 for "Biomolecular condensation of ERC1 recruits ATG8 and NBR1 to drive autophagosome formation for plant heat tolerance"

### Supplemental Figure S1

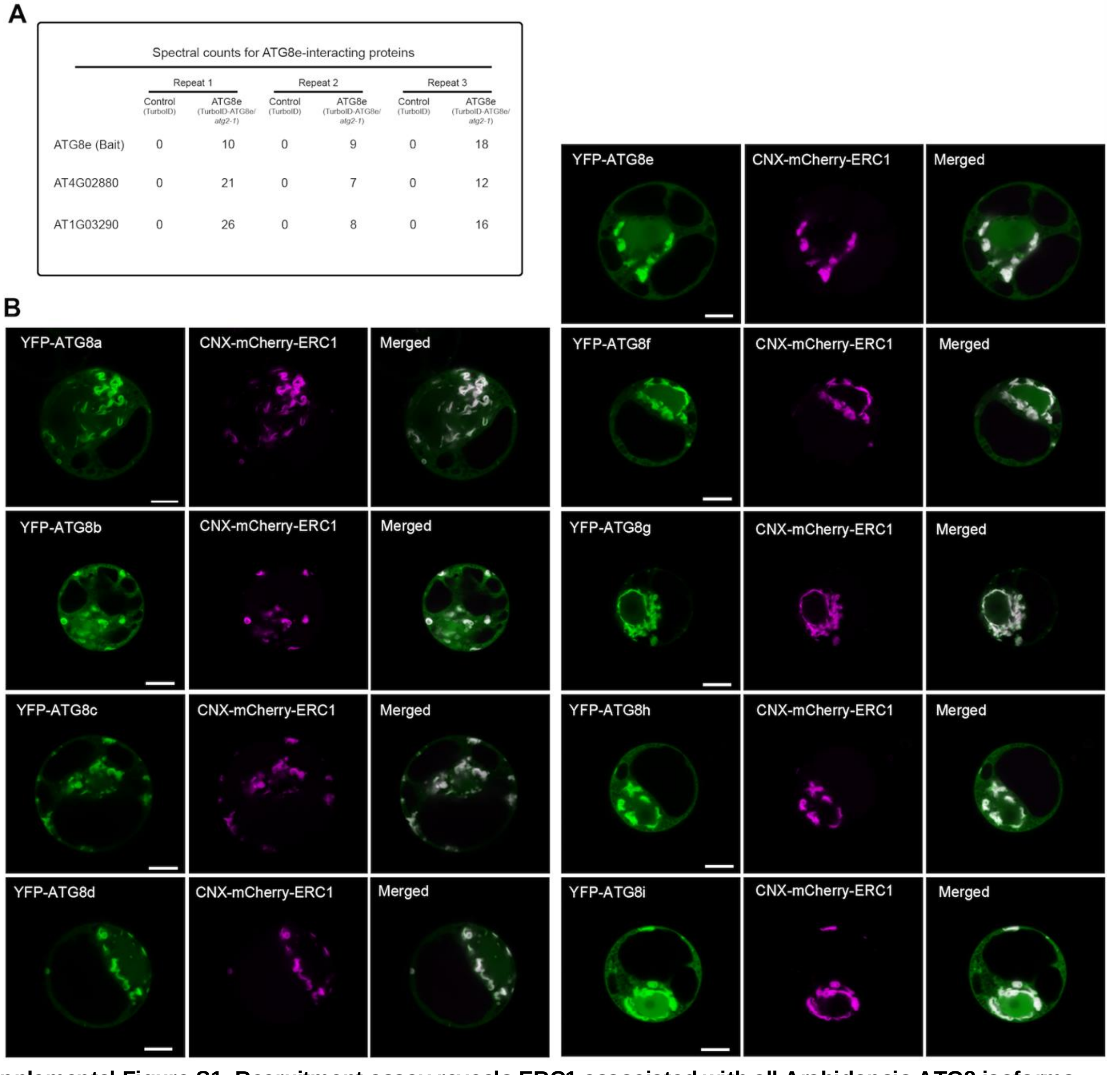

Spectral counts for ATG8e-interacting proteins

| | Repeat 1 | | Repeat 2 | | Repeat 3 | |
|---|---|---|---|---|---|---|
| | Control (TurboID) | ATG8e (TurboID-ATG8e/ *atg2-1*) | Control (TurboID) | ATG8e (TurboID-ATG8e/ *atg2-1*) | Control (TurboID) | ATG8e (TurboID-ATG8e/ *atg2-1*) |
| ATG8e (Bait) | 0 | 10 | 0 | 9 | 0 | 18 |
| AT4G02880 | 0 | 21 | 0 | 7 | 0 | 12 |
| AT1G03290 | 0 | 26 | 0 | 8 | 0 | 16 |



**Supplemental Figure S1. Recruitment assay reveals ERC1 associated with all Arabidopsis ATG8 isoforms.**
(**A**) Proteins identified by tandem mass spectrometry after proximity labeling of TurboID-ATG8e/*atg2-1*. Five-day old transgenic *Arabidopsis* seedlings expressing TurboID or TurboID-ATG8e were subjected to biotin treatment and affinity purification following by LC-MS/MS analysis to identify potential interactors. ERC1(AT4G02880) and ERC2(AT1G03290) were identified as proximity protein of ATG8e. The number of peptides identified in MS of shown.
**(B)** CNX-mCherry-ERC1 and YFP-ATG8s (a-i) were transiently co-expressed in Arabidopsis protoplasts and subjected to confocal observation. Scale bar=10 µm.

### Supplemental Figure S2

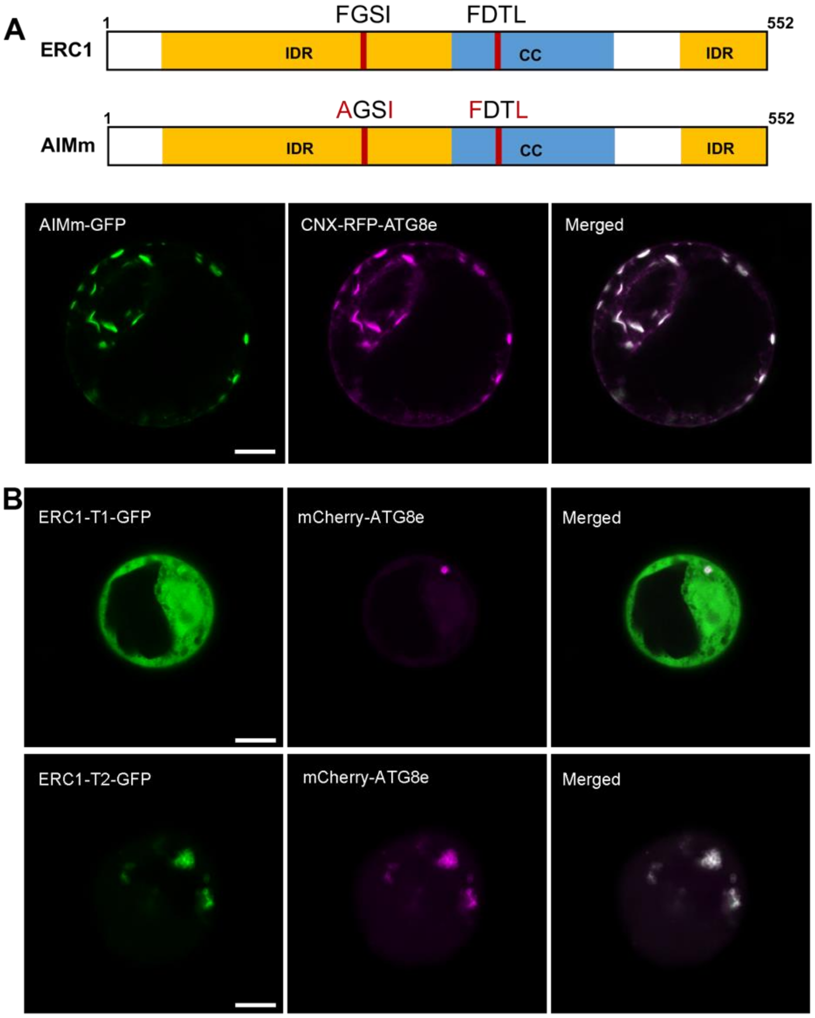


**Supplemental Figure S2. Recruitment assay showed that ERC1-ATG8 association is independent of the predicted AIM motifs.**

**(A)** Two predicted AIM sequences and corresponding site mutations (AIMm) of ERC1 were depicted in the diagram. GFP tagged ERC1 AIMm variant with CNX-RFP-ATG8e were transiently expressed in PSBD protoplasts for 16-18 h before subsequent confocal imaging. Scale bar= 10 µm.

**(B)** Co-expression of ERC1 truncations (T1 and T2) with mCherry-ATG8e in PSBD protoplasts. Constructs were transiently expressed in PSB-D protoplasts for 16-18 h before subsequent confocal imaging. Scale bar: 10 µm.

### Supplemental Figure S3

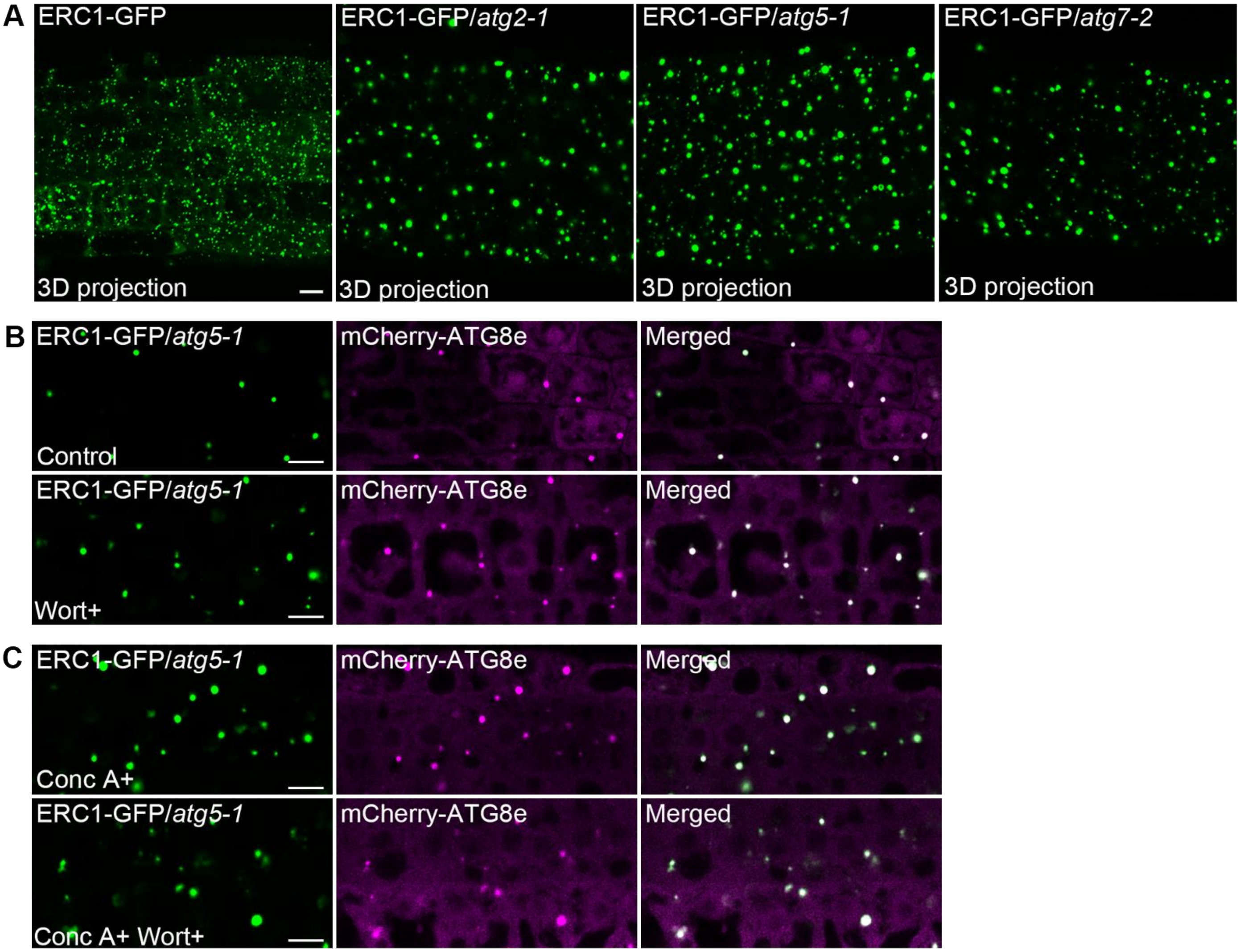


**Supplemental Figure S3. Subcellular analysis of ERC1-GFP/mCherry-ATG8e and drug treatment of ERC1-GFP and ERC1-GFP/*atg5-1* in Arabidopsis root cells.**

**(A**) Subcellular localization of 5-day-old seedlings of ERC1-GFP/*atg2-1*, ERC1-GFP/*atg5-1* or ERC1-GFP/*atg7-2* observed by confocal imaging. Scale bar=10 µm.

**(B)** Large ERC1-GFP and mCherry-ATG8e positive puncta in *atg5-1* mutant plants are insensitive to wortmannin treatment. 5-days-old transgenic seedlings expressing ERC1-GFP x mCherry-ATG8e proteins in *atg5-1* mutant background were incubated in ½ MS medium with or without wortmannin before imaging by confocal microscopy Scale bar=10 µm.

**(C)** Conc A treatment, or combination of Conc A and wortmannin treatment did not interfere the large puncta labeled by ERC1-GFP and mCherry-ATG8e. 5-days-old transgenic seedlings expressing ERC1-GFP x mCherry-ATG8e proteins in *atg5-1* mutant background were incubated in ½ MS medium with Conc A and with or without wortmannin before imaging by confocal microscopy. Scale bar=10 µm.

#### Supplemental Figure S4

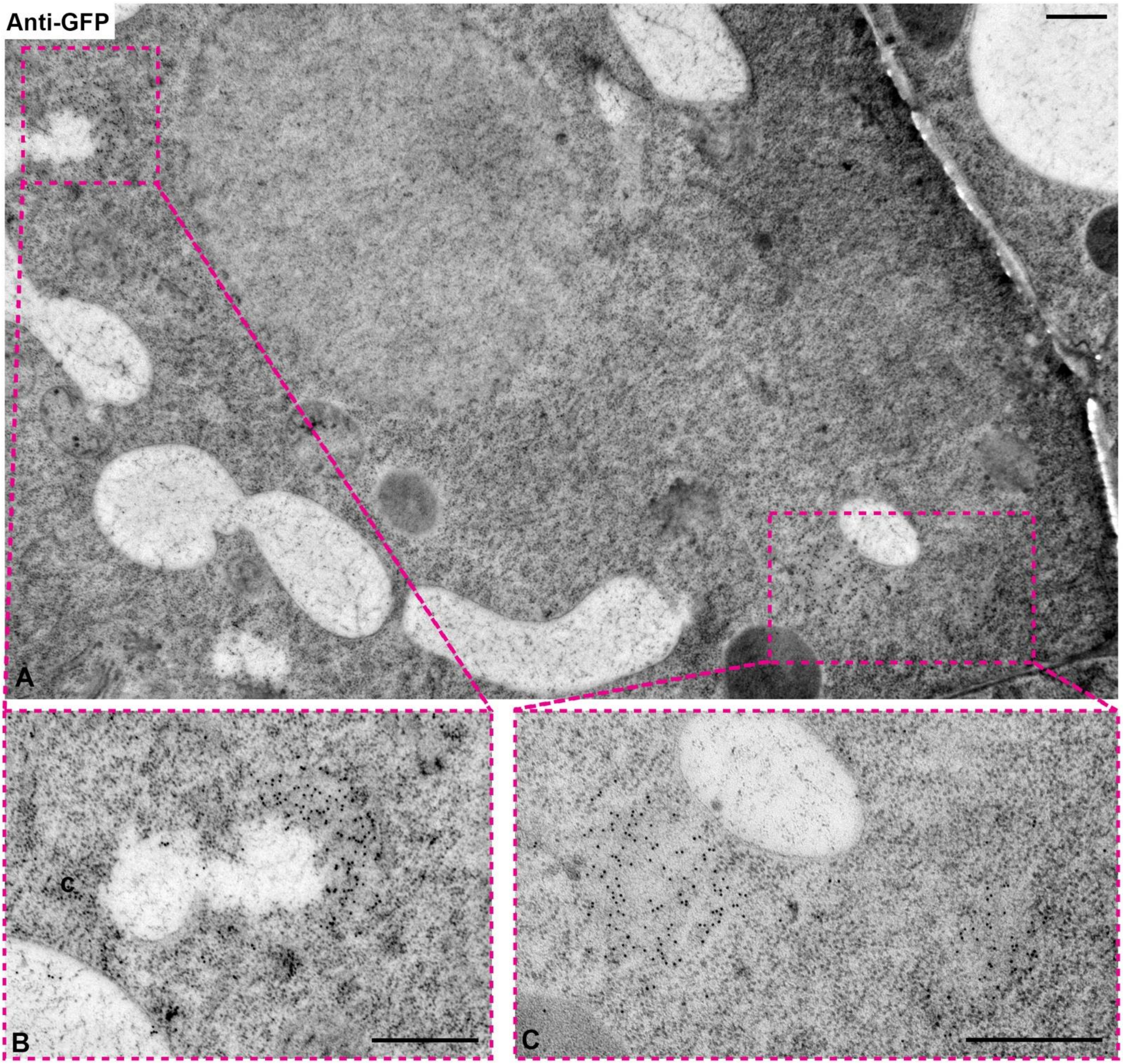


**Supplemental Figure S4. Immuno-EM analysis of ERC1-GFP in root cells after heat stress.** (**A**) A representative example showed membraneless structures (indicated by dashed boxes) with gold particles against anti-GFP. Areas from the indicated dashed boxes were enlarged and shown in **(B)** and **(C).** 5-days-old Arabidopsis seedlings of ERC1-GFP were incubated in preheated liquid 1/2 MS medium under 37 °C for 1 hour, recovered 3 h at 22 °C, followed by high-pressure freezing fixation and sections were immunolabeled with anti-GFP antibodies. Scale bar = 500 nm.

### Supplemental Figure S5

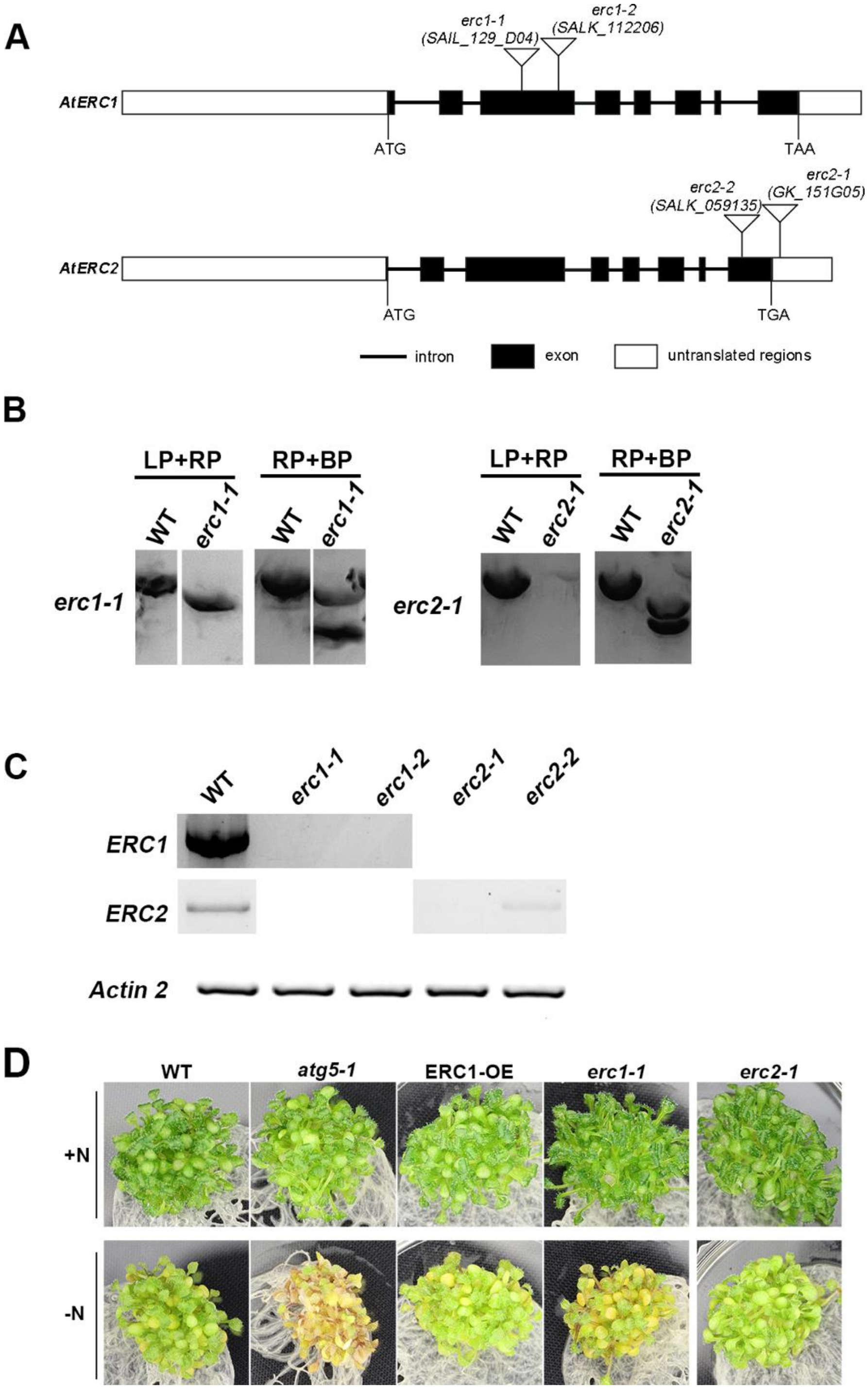


**Supplemental Figure S5. Phenotype analysis of *erc* mutant under starvation condition.**
(A) Schematic diagram showing the T-DNA insertions in the *erc1* and *erc2* mutants respectively.
(B) Genotyping analysis of the *erc1* and *erc2* mutants respectively.
(C) RT-PCR analysis of the *erc1* and *erc2* mutants respectively.
(D) Phenotype analysis under nitrogen starvation conditions. 7-days-old Arabidopsis seedlings were incubated with liquid ½ MS medium (control) or liquid ½ nitrogen-free MS medium. The seedlings were put back to the growth chamber with shaking for an extra 14 days before observation.
